# *DLG2* knockout reveals neurogenic transcriptional programs underlying neuropsychiatric disorders and cognition

**DOI:** 10.1101/2020.01.10.898676

**Authors:** Bret Sanders, Daniel D’Andrea, Mark O. Collins, Elliott Rees, Tom G. J. Steward, Ying Zhu, Gareth Chapman, Sophie E. Legge, Antonio F. Pardiñas, Adrian J. Harwood, William P. Gray, Michael C. O’Donovan, Michael J. Owen, Adam C. Errington, Derek J. Blake, Daniel J. Whitcomb, Andrew J. Pocklington, Eunju Shin

## Abstract

Brain development requires a complex choreography of cell proliferation, specialisation, migration and network formation, guided by the activation and repression of gene expression programs. It remains unclear how this process is disrupted in neuropsychiatric disorders. Here we integrate human genetics with transcriptomic data from the differentiation of human embryonic stem cells into cortical excitatory neurons. This reveals a cascade of transcriptional programs, activated during early corticoneurogenesis *in vitro* and *in vivo*, in which genetic variation is robustly associated with neuropsychiatric disorders and cognitive function. Within these early neurogenic programs, genetic risk is concentrated in loss-of-function intolerant (LoFi) genes, capturing virtually all LoFi disease association. Down-regulation of these programs in *DLG2* knockout lines delays expression of cell-type identity alongside marked deficits in neuronal migration, morphology and action potential generation, validating computational predictions. These data implicate specific cellular pathways and neurodevelopmental processes in the aetiology of multiple neuropsychiatric disorders and cognition.

## Introduction

Although subsequently expanding to encompass other disorders, this study initially sought to explore the role of developmental processes in schizophrenia (SZ). SZ is highly heritable^1, 2^, with genetic variation across the frequency spectrum contributing to disease risk^3–7^. While rare variant studies consistently implicate the disruption of specific mature postsynaptic complexes in SZ aetiology^4, 5, 8–11^, the precise cellular pathways mediating common variant risk (an estimated 30-50% of the total genetic contribution to liability^5^) remain largely obscure. Synaptic complexes enriched for SZ rare variants display little evidence for GWAS association, and while there is convincing evidence for enrichment in broader synapse-related gene sets^12, 13^, these only capture a modest proportion of the overall GWAS signal^12^. In contrast, nearly 50% of genic SNP-based heritability is captured by loss-of-function intolerant (LoFi) genes^12^. Being under extreme selective constraint, LoFi genes are likely to play important developmental roles and are known to be enriched for rare variants contributing to autism spectrum disorders (ASD) and intellectual disability/severe neurodevelopmental delay (ID/NDD)^14^ as well as SZ^11, 15, 16^. This suggests that a significant proportion of SZ common variants may contribute to disease via the disruption of neurodevelopmental pathways harbouring a concentration of LoFi genes, with more severe genetic perturbation of these pathways increasing risk of SZ and disorders associated with more severe developmental phenotypes (e.g. ASD, ID/NDD).

Supporting a neurodevelopmental role for SZ common variants, there is growing evidence that many such risk factors impact gene expression in the foetal brain^17–20^ and are enriched in cell-types at multiple stages of cortical excitatory neuron development^21^. This raises the question: do SZ common variants converge on specific gene expression (transcriptional) programs that are normally activated or repressed during foetal cortical excitatory neuron development? Mutations disrupting key regulators of such programs would be expected to possess a higher contribution to disease risk, reflected in a larger effect size and lower allele frequency. We therefore sought rare, single-gene mutations linked to SZ where the affected gene is expressed in human foetal brain and has the potential to regulate developmental processes. This led us to *DLG2*: multiple independent deletions have been identified at the *DLG2* locus in both SZ and ASD patients^8, 22^; *DLG2* mRNA is present from 8 weeks post-conception in humans^23^ and throughout all stages of *in vitro* differentiation from human embryonic stem cells (hESCs) to cortical projection neurons^24^. Furthermore, the invertebrate orthologue of *DLG1-4* (*Dlg)* is a core component of the *Scrib* signalling module, which regulates cell polarity, differentiation and migration during development^25^. Primarily studied as a post-synaptic scaffold protein, DLG2 is required for the formation of NMDA receptor complexes^26^: these complexes regulate the induction of several forms of synaptic plasticity^27^ and are enriched for rare mutations in SZ cases^4, 5, 8–11^. This raises the intriguing possibility that DLG2 may be required for the normal operation of both adult and developmental signalling pathways relevant to SZ pathophysiology.

To explore the role of DLG2 in neurodevelopment we engineered homozygous loss-of-function *DLG2* mutations into hESCs using the CRISPR-CAS9 system. Mutant (*DLG2^-/-^*) and isogenic sister wild-type (WT) hESC lines were differentiated into cortical excitatory neurons and cells characterised at multiple developmental timepoints to identify phenotypes and gene expression changes in *DLG2^-/-^* lines (Fig. 1a). Neurodevelopmental expression programs dysregulated in *DLG2^-/-^* lines were identified and analysed for risk variant enrichment; we then explored the biological function of disease-relevant programs, both computationally and experimentally, and evaluated the contribution of LoFi genes to common and rare variant associations (Fig. 1a).

**Figure 1.**
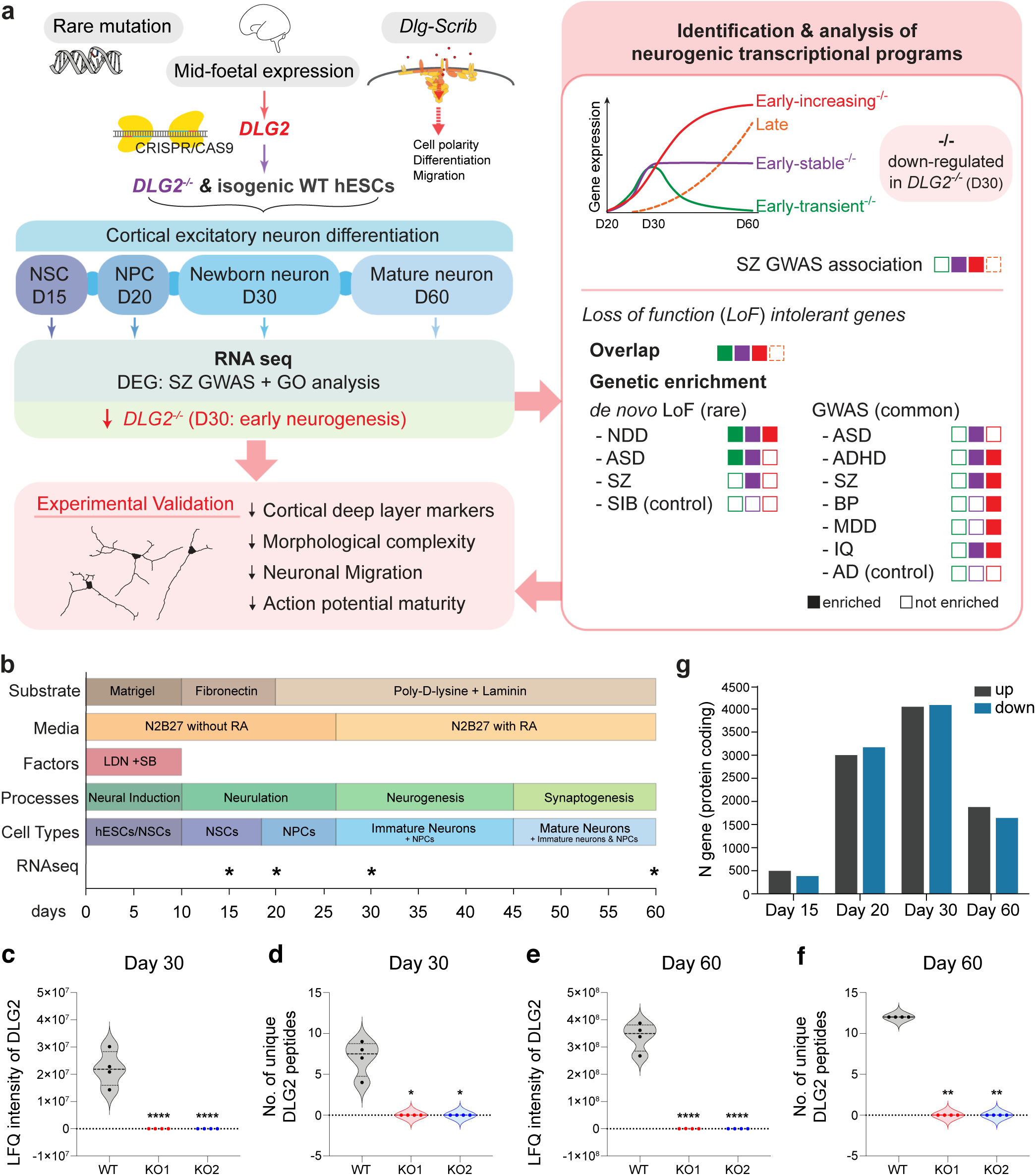
Study design and number of differentially expressed genes. **a**, Study summary. *DLG2^-/-^* hESCs were generated via the CRISPR/CAS9 system; a sister hESC line that went through the gene editing protocol but remained as WT served as a control. hESCs were differentiated into cortical excitatory neurons and RNA collected at multiple timepoints: predominant cell types shown for each timepoint. Genetic analysis revealed SZ GWAS enrichment in genes down-regulated at day 30 in *DLG2^-/-^* lines, coinciding with early neurogenesis: corresponding phenotypes predicted via GO term analysis were validated experimentally. Transcriptional programs active during neurogenesis were identified based on differential WT gene expression across successive timepoints. SZ common variant risk was concentrated in two early neurogenic programs down-regulated in *DLG2^-/-^* cells. Loss of function intolerant (LoFi) genes – highly over-represented in all 3 early programs – were enriched for common and rare (*de novo* LoF) variants contributing to a wide range of neuropsychiatric disorders and cognition; no enrichment was seen in *de novo* LoF mutations from unaffected siblings of ASD cases or common risk variants for Alzheimer’s disease. **b**, Overview of cortical differentiation protocol with approximate timings of key developmental processes and predominant cell types present in culture. Asterisks indicate timepoints selected for RNA sequencing. **c-f**, Label free quantification (LFQ) of DLG2 protein levels in PDZ-ligand (NR2 C-terminus) affinity pulldowns in day 30 and 60 WT and *DLG2^-/-^* cells using LC-MS/MS analysis confirms the absence of PDZ-ligand binding DLG2 in knockout cells. One-way ANOVA with bonferroni multiple comparison correction applied to **c** (F_2,9_=45.54, P<0.0001) and **e** (F_2,9_=172.9, P<0.0001). Kruskal-Wallis test with Dunn’s multiple comparison correction applied to **d** (H(2)=10.46, p=0.0061) and **f** (H(2)=11.00, p=0.0061). *p<0.05; **p<0.01; ****p<0.0001 vs. WT. **g**, Number of protein coding genes differentially expressed in *DLG2^-/-^* cells relative to WT at each timepoint.

## Results

### Knockout generation and validation

Two *DLG2^-/-^* lines were created from H7 hESCs using the CRISPR/Cas9-D10A nickase system targeting the first PDZ domain (Supplementary Fig. 1). Sequencing of predicted off-target sites revealed no mutations (Methods, Supplementary Fig. 2 & Supplementary Table 1). All subsequent analyses compared these lines to an isogenic WT sister line that went through the same procedure but remained genetically unaltered.

*DLG2^-/-^* and WT lines were differentiated into cortical excitatory neurons using a modified dual SMAD inhibition protocol^28, 29^; RNA was extracted in triplicate from each line at 4 timepoints spanning cortical excitatory neuron development and gene expression quantified (Fig. 1b, Supplementary Fig. 3). A significant decrease in *DLG2* mRNA was observed for exons spanning the first PDZ domain, with a similar decrease inferred for PDZ-containing transcripts, indicating degradation of *DLG2^-/-^* transcripts via nonsense-mediated decay (Supplementary Fig. 4). Quantitative mass spectrometry-based proteomic analysis of peptide-affinity pulldowns using the NMDA receptor NR2 subunit PDZ peptide ligand^30^ confirmed the presence of DLG2 in pulldowns from WT but not *DLG2^-/-^* lines (Fig. 1c-f, Supplementary Table 2). Genotyping revealed no CNVs in either *DLG2^-/-^* line relative to WT (Supplementary Fig. 5a). Both *DLG2^-/-^* lines expressed pluripotency markers OCT4, SOX2 and NANOG at 100% of WT levels (Supplementary Fig. 5b). Cells were extensively characterised for their cortical identity using western blotting and immunocytochemistry from days 20-60. Over 90% of day 20 cells were positive for FOXG1, PAX6 and SOX2 and <1% cells expressed ventral genes such as DLX1, GBX2, NKX2.1 and OLIG3 (Supplementary Fig. 6), confirming dorsal forebrain fate. In addition, staining of markers expressed in ventral forebrain-derived neurons from striatal, thalamic and hypothalamic nuclei confirmed no or trace expression (Supplementary Fig. 6).

### DLG2 knockout impacts gene expression during cortical excitatory neuron development

To robustly identify genes dysregulated by *DLG2* knockout, expression data from the two *DLG2^-/-^* lines was pooled and compared to WT at each timepoint (Methods). Disruption of *DLG2* had a profound effect: of the >13,000 protein-coding genes expressed at each timepoint, ∼7% were differentially expressed between *DLG2^-/-^* and WT at day 15, rising to 40-60% between days 20 and 30 then decreasing to ∼25% by day 60 (Fig. 1g, Supplementary Table 3).

### Common risk variants implicate disruption of neurogenesis in SZ

We next tested whether genes differentially expressed in *DLG2^-/-^* lines at each timepoint were enriched for SZ common risk variants. Taking summary statistics from the largest available SZ GWAS^12^, we utilised the competitive gene-set enrichment test implemented in MAGMA (version 1.07)^31^. As expected, the set of all genes expressed at one or more timepoint in *DLG2^-/-^* or WT lines (*all^WT+KO^*) was highly enriched for common variant association (P = 2.1 x 10^-17^) reflecting the neural lineage of these cells. We therefore tested genes up- and down-regulated at each timepoint for genetic association conditioning on *all^WT+KO^* using the strict *condition-residualize* procedure (all subsequent GWAS enrichment tests were conditioned on *all^WT+KO^* in the same way). This revealed strong association enrichment solely in genes down-regulated at day 30 (30_down_^-/-^: P_corrected_ = 1.9 x 10^-7^, Fig. 2a), coinciding with active neurogenesis (Fig. 1b). When compared to *all^WT+KO^*, 30_down_^-/-^ genes were over-represented in GO terms related to neuronal development, function and migration (Methods, Supplementary Table 4). Iterative refinement via conditional analyses identified 23 terms with independent evidence for over-representation (Fig. 2b, Methods). This suggests that loss of *DLG2* dysregulates transcriptional programs underlying neurogenesis (neuronal growth, electrophysiological properties and migration) and implicates these processes in SZ aetiology.

**Figure 2.**
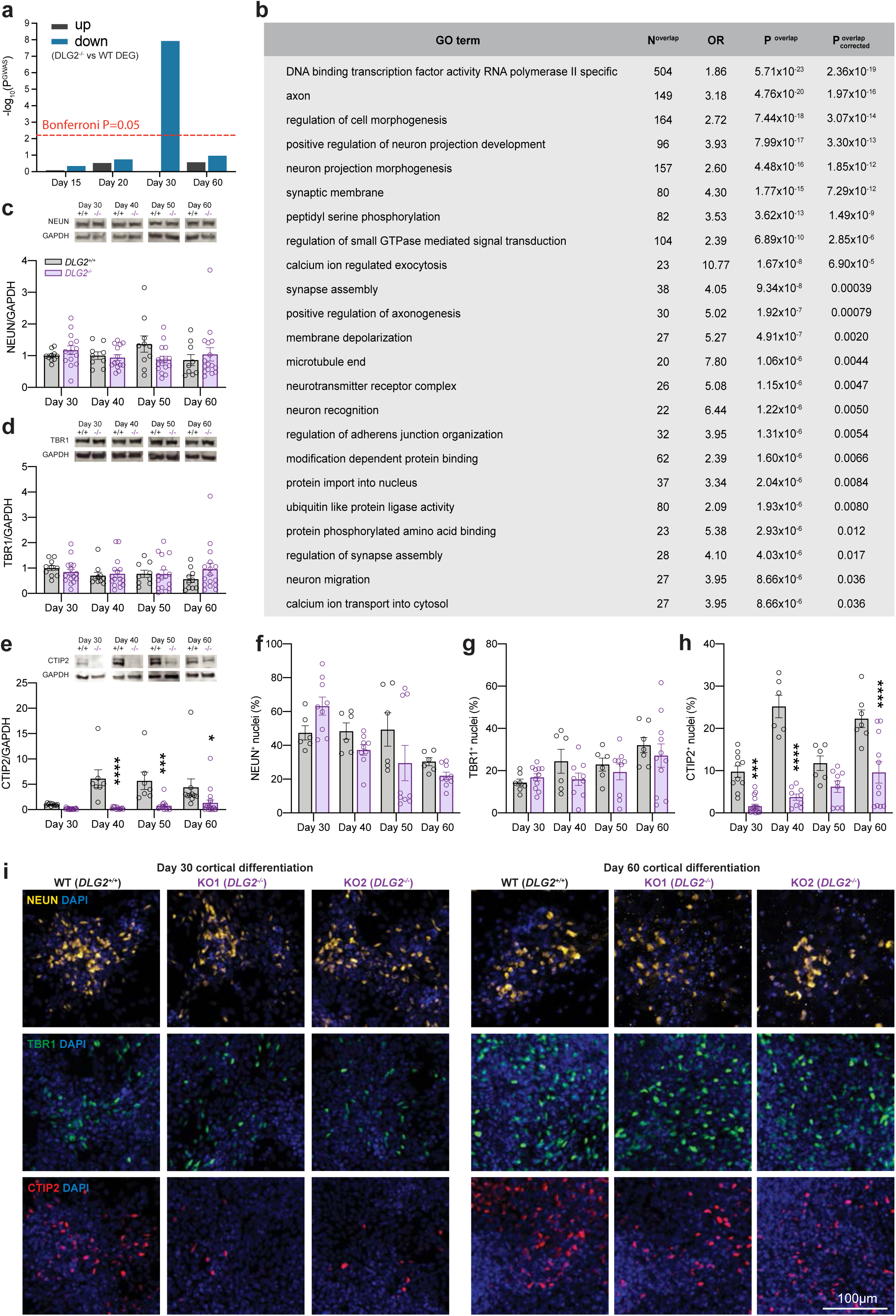
Common risk variants implicate disruption of neurogenesis in schizophrenia. **a**, Enrichment for common schizophrenia risk variants in genes up- & down-regulated at each timepoint (*DLG2^-/-^* relative to WT), conditioning on all expressed genes. **b**, Gene ontology (GO) terms over-represented amongst genes down-regulated at day 30 in *DLG2^-/-^* lines relative to all expressed genes. **c**, NEUN western blot protein quantification. Neither genotype (F_1,90_=0.1852; P=0.6680; n≥7) nor time (F_3,90_=0.5382; P=0.6573; n≥7) had significant effects on NEUN expression. **d**, TBR1 western blot protein quantification. Neither genotype (F_1,95_=0.3899; P=0.5338; n≥9) nor time (F_3,35_=0.5052; P=0.6793; n≥9) had significant effects on TBR1 expression. **e**, CTIP2 western blot protein quantification. Both genotype (F_1,86_=39.89; P<0.0001; n≥7) and time (F_3,86_=5.262; P=0.0022; n≥7) had significant effects on CTIP2 expression. **f**, ICC quantification of NEUN^+^ cells. Time (F_3,52_=7.018, P=0.0005; n≥6) had a significant effect on NEUN expression, while genotype (F_1,52_=1.687; P=0.1998; n≥6) did not. **g**, ICC quantification of TBR1^+^ cells. Time (F_3,58_=4.738, P=0.0050; n≥6) did have a significant effect on TBR1 expression, while genotype (F_1,58_=1.664; P=0.2022; n≥6) did not. **h**, ICC quantification of CTIP2^+^ cells. Both genotype (F_1,67_=101.8; P<0.0001; n≥6) and time (F_3,67_=18.93; P<0.0001; n≥6) had significant effects on CTIP2 expression. **i**, Representative ICC images of NEUN, TBR1 and CTIP2 with DAPI nuclear counterstain. Western blotting (**c**-**e**) and ICC data sets (**f**-**h**) were analysed by two-way ANOVA with post hoc comparisons using Bonferroni correction, comparing to WT controls. Stars above bars represent Bonferroni-corrected post hoc tests, *P<0.05; **P<0.01; ***P<0.001; ****P<0.0001 vs. WT control. All data presented as mean ± SEM.

### Dysregulation of neurogenesis in DLG2^-/-^ lines delays cortical cell-fate expression

To validate disruption of neurogenesis in *DLG2^-/-^* lines and investigate whether this leads to differences in the number or type of neurons produced, we compared the expression of cell-type specific markers in *DLG2^-/-^* and WT lines from days 30-60 via immunocytochemistry (ICC) and Western blotting (Fig. 2c-i). From ICC it was clear that *DLG2^-/-^* cells are able to differentiate and produce postmitotic neurons expressing characteristic neuronal markers such as NEUN and TUJ1 plus cortical deep layer markers TBR1 and CTIP2 (Fig. 2c-i, Supplementary Fig. 7). Western blot of NEUN (Fig. 2c) and MAP2 (Supplementary Fig. 7) and quantification of NEUN^+^ cells following ICC (Fig. 2f) revealed no difference in the percentage of neurons produced by *DLG2^-/-^* cultures. This is in line with the comparable percentage of cells in the cell cycle/neural progenitors at days 30-60 in *DLG2^-/-^* and WT cultures indicated by a similar proportion of KI67^+^ and SOX2^+^ cells (Supplementary Fig. 7). At these early timepoints we would not expect to see the generation of upper layer neurons. Although we could identify a small percentage of SATB2^+^ cells in both WT and KO lines, all co-expressed CTIP2 (Supplementary Fig. 7) indicating their deep layer identity^32^. An analysis of deep layer markers TBR1 and CTIP2 revealed a significant decrease in CTIP2^+^ cells but a comparable proportion of TBR1^+^ neurons for all timepoints investigated (Fig. 2d, e, g-i). On average the proportion of CTIP2^+^ cells recovered from 15% of the WT level on day 30 to 50% by day 60, although there was notable variation between *DLG2^-/-^* lines (Supplementary Fig. 8); total CTIP2 protein level also recovered to some extent, but at a slower rate (Supplementary Fig. 8). Thus, *DLG2^-/-^* does not affect the rate at which neurons are produced but delays the expression of subtype identity in new-born deep layer neurons.

### DLG2^-/-^ lines display deficits in neuron morphology & migration

Given the over-representation of 30_down_^-/-^ genes in terms related to neuron morphogenesis and migration (Fig. 2b), we sought to experimentally validate these phenotypes. Immature (day 30) and mature (day 70) neurons were traced and their morphology quantified (Fig. 3). At both timepoints *DLG2^-/-^* neurons displayed a simpler structure than WT, characterised by a similar number of primary neurites projecting from the soma (Fig. 3a) but with greatly reduced branching (Fig. 3b). Total neurite length did not differ (Fig. 3c), leading to a clear *DLG2^-/-^* phenotype of longer and relatively unbranched primary neurites (Fig. 3e). There was no significant difference in soma area (Fig. 3d). Day 40 *DLG2^-/-^* neurons had a slower speed of migration (Fig. 3f) and reduced displacement from their origin after 70 hrs (Fig. 3g, h). In summary, *DLG2^-/-^* neurons show clear abnormalities in both morphology and migration, validating the GO term analysis.

**Figure 3.**
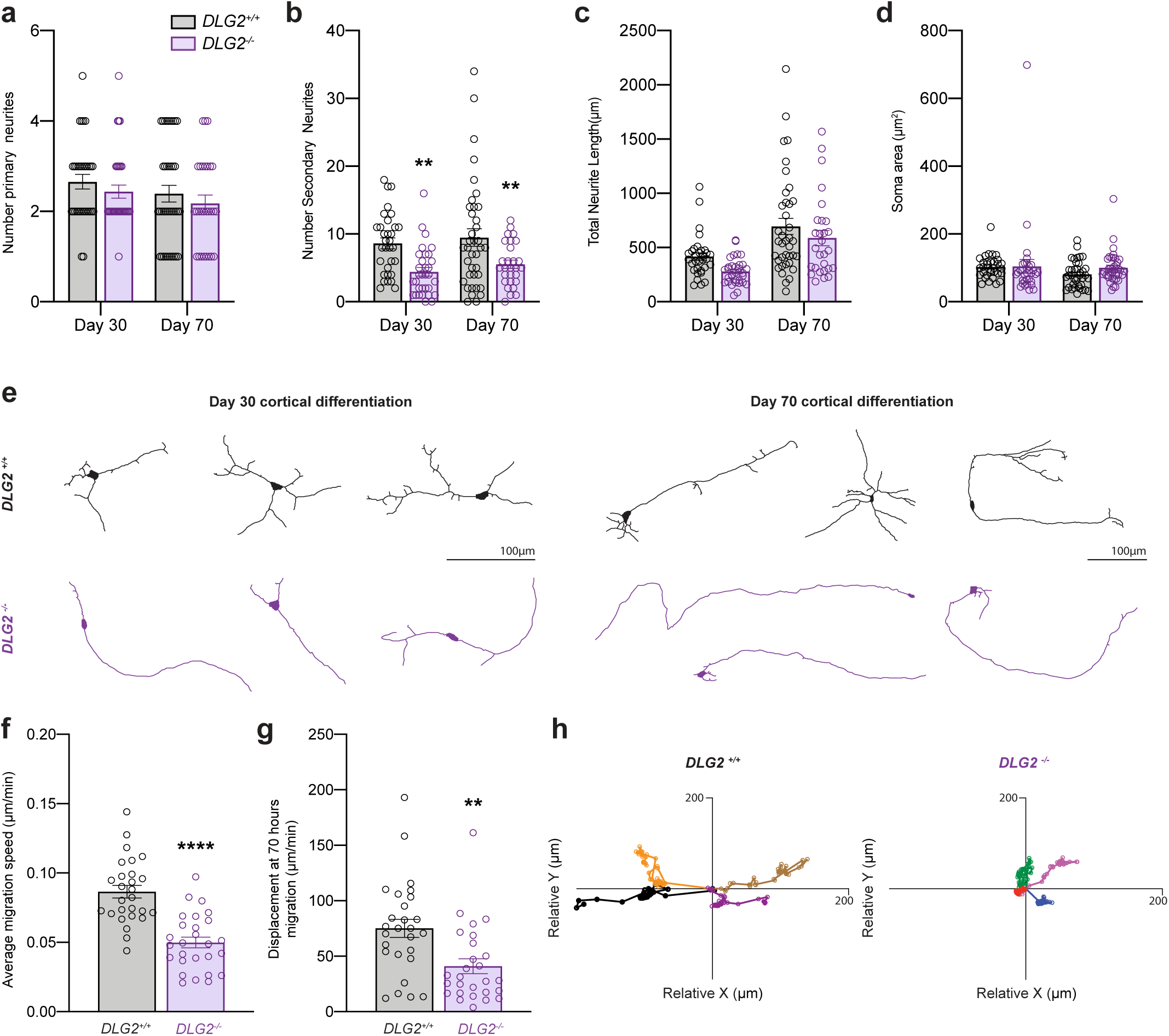
*DLG2^-/-^* lines display deficits in neuron morphology & migration. **a**, The number of primary neurites (projecting from the soma). Neither genotype (F_1,126_=1.591; P=0.2095; n≥28) nor time (F_1,126_=2.278; P=0.1337; n≥28) had significant effects the numbers of primary neurites. **b**, The number of secondary neurites (projecting from primary neurites). Genotype (F_1,126_=18.78, P<0.0001; n≥28) had a significant effect on number of secondary neurites, while time (F_1,126_=1.082, P=0.3003; n≥28) did not. **c**, The total neurite length. Both genotype (F_1,126_=4.568; P=0.0345; n≥28) and time (F_1,126_=26.33; P<0.0001; n≥28) had significant effects on total neurite length. However, post hoc analysis showed no significant differences at individual timepoints. **d**, The soma area. Neither genotype (F_1,136_=; P=0.9170; n≥28) nor time (F_1,136_=1.399; P=0.2390; n≥28) had a significant effect on soma area. **e**, Representative traces showing the neuronal morphology. **f**, The average speed of neuronal migration over 70 hours, from day 40 of cortical differentiation. *DLG2^-/^*^-^ neurons showed significantly decreased average migration speed compared to WT (t_52_=6.1175; P<0.0001; n=27). **g**, The displacement of neurons at 70 hours migration. *DLG2^-/^*^-^ neurons showed significantly decreased displacement compared to WT (t_52_=3.244; P=0.0021; n=27). **h**, Representative traces of neuronal migration from a given origin over 70 hours. Morphology data sets (**a**-**d**) were analysed by two-way ANOVA with post hoc comparisons using Bonferroni correction, comparing to WT controls. Migration data sets (**f**,**g**) were analysed by unpaired two-tailed Student’s t-test. Stars above bars represent, **P<0.01; ****P<0.0001 vs. WT control (Bonferroni-corrected for morphology analyses). All data presented as mean ± SEM.

### Distinct transcriptional programs regulated by DLG2 are enriched for common SZ risk alleles

We postulated that loss of *DLG2* inhibits the activation of transcriptional programs driving neurogenesis, which starts between days 20 and 30 and steadily increases thereafter. If this is the case, then SZ genetic enrichment in 30_down_^-/-^ should be captured by genes normally upregulated between days 20 and 30 in WT cultures (20-30_up_^WT^). Analysing differential expression between WT samples at successive timepoints, we found strong risk variant enrichment in 20-30_up_^WT^ (Fig. 4a). The overlap between 20-30_up_^WT^ and 30_down_^-/-^ captured the signal in both sets (P_overlap_ = 3.23 x 10^-10^; 30_down_^-/-^ only P = 0.44; 20-30_up_^WT^ only P = 0.62). This was not simply due to the size of the overlap (3075 genes, 85% of 20-30_up_^WT^) as the regression coefficient for the set of overlapping genes (β = 0.14), which reflects magnitude of enrichment, was significantly greater than for genes unique to 30_down_^-/-^ (β = 0.006, P_different_ = 0.0015) or 20-30_up_^WT^ (β = −0.015, P_different_ = 0.0045). Thus, it is neurogenic transcriptional programs that are typically upregulated in WT but down-regulated in *DLG2^-/-^* lines that are enriched for SZ common variants.

**Figure 4.**
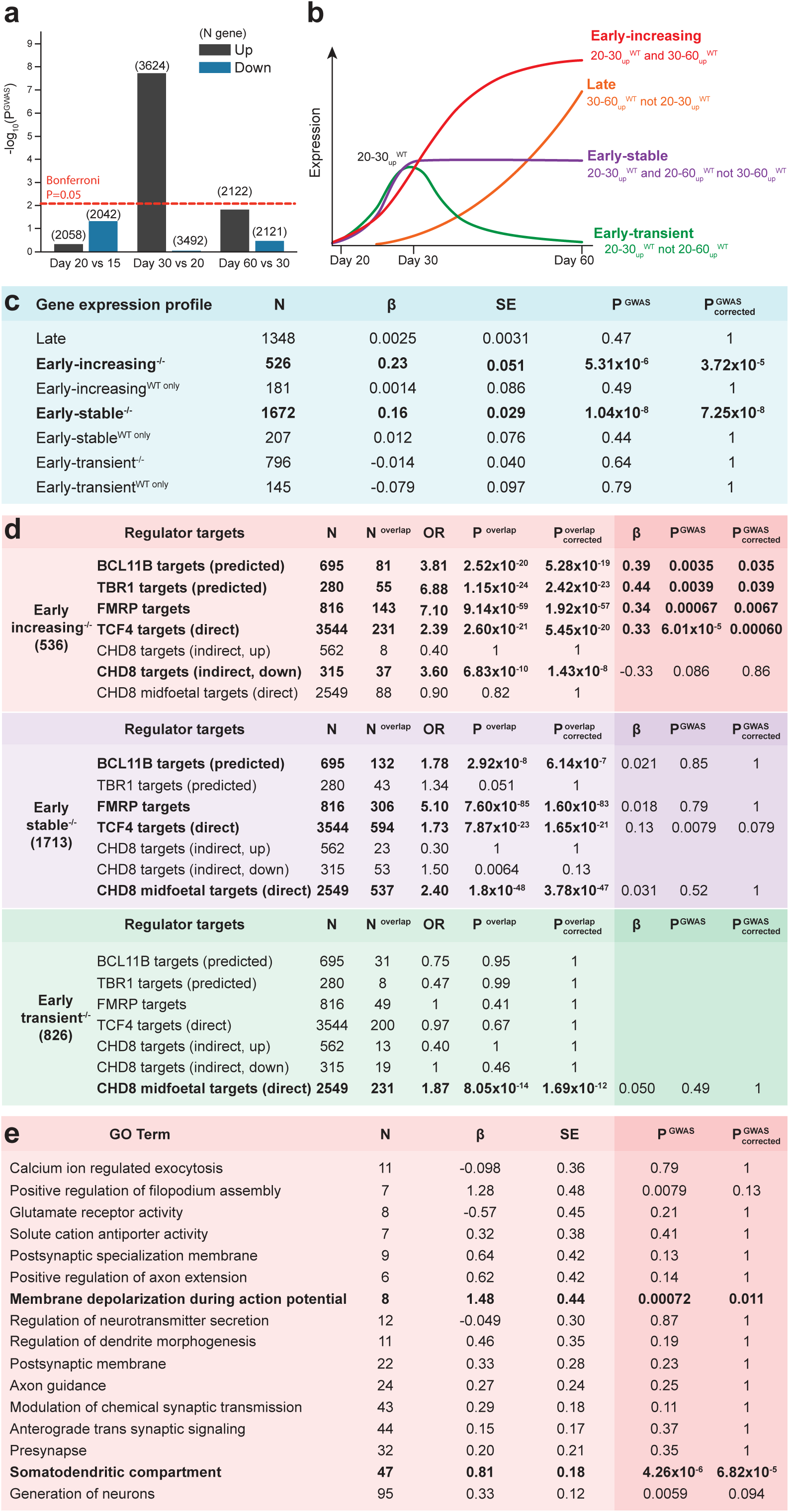
*DLG2* regulates a cascade of transcriptional programs driving neurogenesis & differentiation. **a**, Enrichment for common schizophrenia risk variants in genes up- & down-regulated between each successive pair of timepoints in WT, conditioning on all expressed genes. Dotted line indicates P_corrected_ = 0.05 following Bonferroni correction for 6 tests. **b**, Four discrete transcriptional programs initiated following the onset of neurogenesis were identified based upon WT differential expression between timepoints: *early-increasing*, genes significantly upregulated between days 20 and 30 (20-30_up_^WT^) and also days 30 and 60 (30-60_up_^WT^); *early-stable* genes, present in 20-30_up_^WT^ and 20-60_up_^WT^ but not 30-60_up_^WT^; *early-transient* (20-30_up_^WT^ but not 20-60_up_^WT^); and *late* (30-60_up_^WT^ but not 20-30_up_^WT^). **c**, SZ GWAS enrichment in each transcriptional program, further split into genes that are down-regulated in DLG2^-/-^ lines at day 30 (e.g. early-stable^-/-^) and those that are not (e.g. early-stable^WT only^). Tests condition on all expressed genes. **d**, Identification of programs over-represented for the targets of key regulators (P^overlap^) when compared to all expressed genes; all program-regulator enrichments identified were taken forward for genetic analysis, testing whether regulator targets were more highly enriched for SZ association than other genes in that program (P^GWAS^). **e**, SZ GWAS enrichment in GO terms over-represented amongst early-increasing^-/-^ genes, conditioning on all expressed and all early-increasing^-/-^ genes.

To more precisely identify SZ-relevant transcriptional programs active during neurogenesis, we classified 20-30_up_^WT^ genes based on their subsequent WT expression profiles (Fig. 4b, Methods): early-increasing genes, whose expression continues to rise between days 30 and 60; early-stable genes, whose expression stays at a relatively constant level; and early-transient genes, whose expression is later down-regulated. We also defined a set of late genes, whose expression only increases significantly after day 30. These were further partitioned into genes that were down-regulated at day 30 in *DLG2^-/-^* lines (e.g. early-stable^-/-^) and those that were not (e.g. early-stable^WT only^). The sole exception to this was the late set, which had minimal overlap with 30_down_^-/-^ (62 out of 1399 genes) and was therefore left intact. Early-stable^-/-^ and early-increasing^-/-^ sets were robustly enriched for SZ association (conditioning on *all^WT+KO^*, Fig. 4c), revealing that SZ GWAS association is restricted to 2 distinct transcriptional programs normally activated during the onset of neurogenesis but down-regulated in *DLG2^-/-^* lines.

### Cascade of transcriptional programs predicted to drive neurogenesis & differentiation

We next investigated the biological function of early neurogenic programs dysregulated in *DLG2^-/-^* lines. Each was over-represented for a coherent set of GO terms indicating a distinct biological role (Supplementary Table 5): early-transient^-/-^ for histone/chromatin binding and transcriptional regulation; early-stable^-/-^ for signal transduction, transcriptional regulation, neurogenesis, cell projection development, migration and differentiation; and early-increasing^-/-^ for axon guidance, dendrite morphology, components of pre- and post-synaptic compartments and electrophysiological properties. These functions suggest a linked, time-ordered cascade of transcriptional programs spanning early neurogenesis. This begins with an initial phase of chromatin remodelling (early-transient^-/-^) that establishes neuron sub-type identity and leads to activation of a longer-term program guiding the growth and migration of new-born neurons (early-stable^-/-^). This in turn promotes the fine-tuning of sub-type specific neuronal structure, function and connectivity as cells enter the terminal phase of differentiation (early-increasing^-/-^).

To test support for the existence of this transcriptional cascade and its disruption in disease, we identified disease-relevant regulatory genes from each program whose downstream targets have been experimentally identified or computationally predicted (Methods). Reflecting our hypothesis that dysregulation of these pathways is likely to play a role in multiple neurodevelopmental disorders, we sought regulators linked to SZ, ASD and ID/NDD: chromatin modifier *CHD8*^33–37^ from early-transient; transcription factor *TCF4* ^12, 38–40^ and translational regulator FMRP^12, 41, 42^ from early-stable^-/-^; and transcription factors (and deep layer markers) *TBR1*^42, 43^ and *BCL11B* (*CTIP2*) ^12, 13, 42, 44^ from early-increasing^-/-^. We predicted that a substantial proportion of early-stable^-/-^ genes would be directly regulated by *CHD8*, while early-increasing^-/-^ would be enriched for genes directly regulated by *TCF4*, *TBR1* and *BCL11B*. Since early-transient^-/-^ is responsible for activating subsequent programs, we predicted that early-increasing^-/-^ would be enriched for indirect targets of *CHD8* (genes not directly regulated but whose expression is altered when *CHD8* is perturbed) that are down-regulated in *CHD8* knockdown cells^36^. We also predicted that early-transient^-/-^ genes would *not* be enriched for targets of terminal phase regulators *BCL11B* and *TBR1*. FMRP represses the translation of its mRNA targets, facilitating their translocation to distal sites of protein synthesis^41, 45^, and its function is known to be important for axon and dendrite growth^46^. We therefore predicted that early-stable^-/-^ and -increasing^-/-^ (but not -transient^-/-^) genes would be enriched for FMRP targets. Over-representation tests emphatically confirmed these predictions, supporting the existence of a regulatory cascade driving early neurogenic transcriptional programs disrupted in neuropsychiatric disorders (Fig. 4d). In addition, the targets of *TCF4*, FMRP, *BCL11B* and *TBR1* were more highly enriched for SZ association than other genes in early-increasing^-/-^ (Fig. 4d), highlighting specific pathways through which these known risk genes are likely to contribute to disease.

### Convergence of genetic risk on perturbed action potential generation

We next tested whether biological processes over-represented in early-stable^-/-^ or early-increasing^-/-^ (Supplementary Table 5) captured more or less of the SZ association in these programs than expected (Methods). None of the 13 semi-independent GO term subsets identified in early-stable^-/-^ differed substantially from early-stable^-/-^ as a whole (Supplementary Table 6), indicating that risk factors are distributed relatively evenly between them. Of the 16 subsets for early-increasing^-/-^, *somatodendritic compartment* and *membrane depolarization during action potential* displayed evidence for excess enrichment relative to the program as a whole (Fig. 4e). No single term showed evidence for depletion, suggesting that diverse biological processes regulating neuronal growth, morphology and function are perturbed in SZ. The enhanced enrichment in action potential (AP) related genes is particularly striking: while postsynaptic complexes regulating synaptic plasticity are robustly implicated in SZ^4, 5, 8–11^, this represents the first evidence that the molecular machinery underlying AP generation is also disrupted. We therefore sought to confirm the disruption of APs in *DLG2^-/-^* lines (Fig. 5a-j), also investigating the impact of *DLG2* loss on synaptic transmission (Fig. 5l-n).

**Figure 5.**
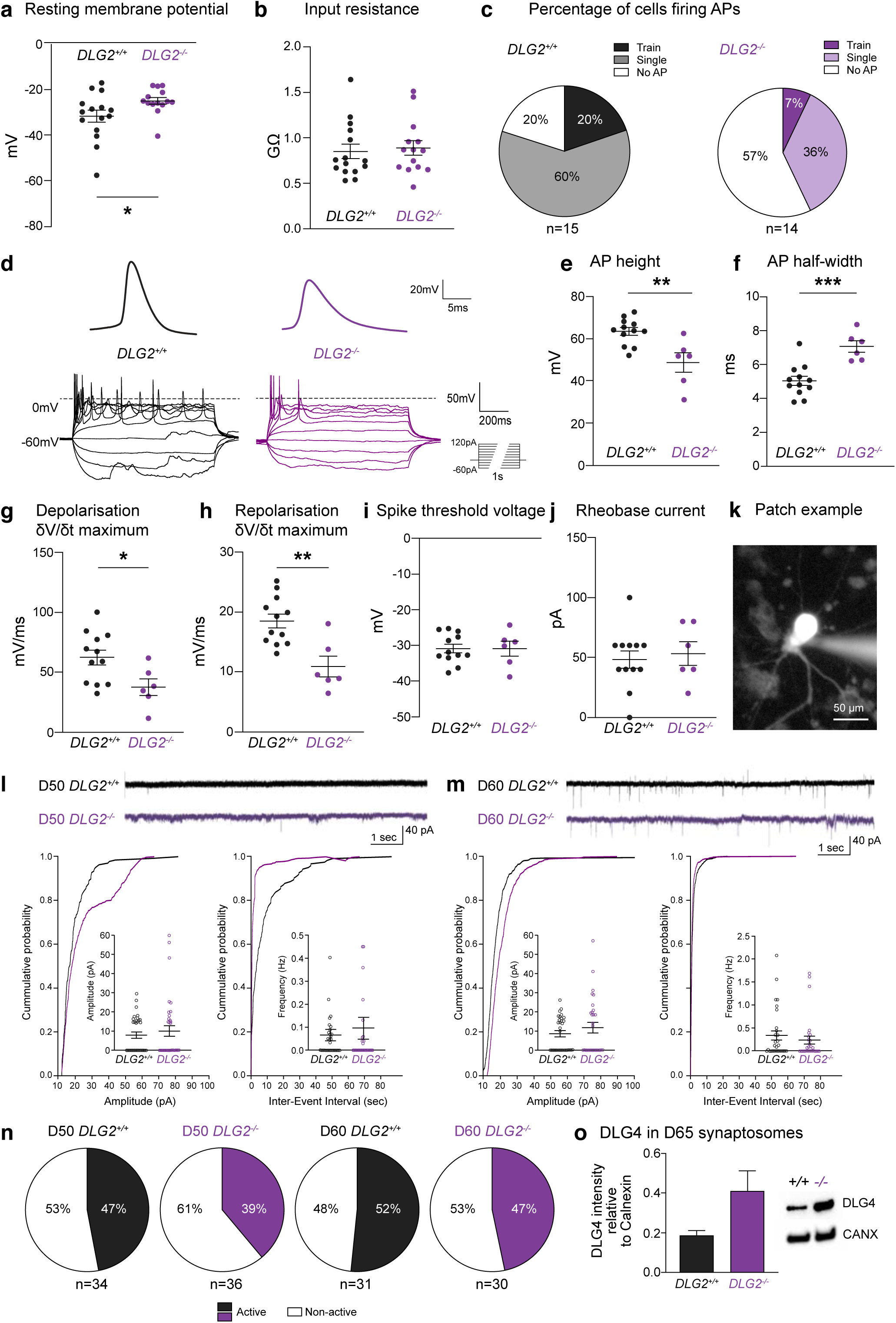
Electrophysiological properties of *DLG2^-/-^* neurons. **a**, Resting membrane potential (t_27_=2.151, P=0.0406) and **b**, input resistance (t_27_=0.3366, P=0.7390) of day 50 WT and *DLG2^-/-^* neurons (n=15 and 14, respectively). **c**, Percentages of cells firing action potentials (APs) upon current step injection. **d**, Example traces of first overshooting AP and APs evoked by current step injection (−60pA to 120pA, increment 20pA, duration 1s). **e**, AP height (t_16_=3.661, P=0.0021), **f**, AP half-width (t_16_=4.462, P=0.0004), **g**, AP maximum depolarising speed (t_16_=2.463, P=0.0255), **h**, AP maximum repolarising speed (t_16_=3.728, P=0.0018), **i**, spike threshold voltage (t_16_=0.004093, P=0.9968) and **j**, rheobase current (t_16_=0.4061, P=0.6900) of day 50 WT and *DLG2^-/-^* neurons (n=12 and 6, respectively) are shown. **k**, Example of day 50 neuron being whole-cell patch clamped with fluorescent dye injection. Spontaneous excitatory postsynaptic current (sEPSC) examples from day 50 (**l**) and 60 (**m**) neurons. Both the amplitude and frequency of day 50 and 60 neurons from WT and *DLG2^-/-^* neurons were comparable (day 50, amplitude: t_68_=0.6974, P=0.4879, frequency: t_66_=0.5467, P=0.5865; day 60 amplitude: t_59_=1.021, P=0.3114, frequency: t_58_=0.7671, P=0.4464). **n**, Percentage of cells displaying sEPSCs. **o**, Western blot analysis of DLG4 in synaptosomes of day 65 WT and *DLG2^-/-^* neurons, displaying trend towards increased DLG4 expression in *DLG2^-/-^* neurons (t_2_=2.157, P=0.1637). *p<0.05; **p<0.01; ***, p<0.001. All data presented as mean ± SEM.

Day 50 *DLG2^-/-^* neurons were less excitable, with a significantly more depolarised resting membrane potential (Fig. 5a). Stepped current injection evoked AP firing in 80% WT but only 43% *DLG2^-/-^* neurons (Fig. 5c). APs produced by *DLG2^-/-^* cells were characteristic of less mature neurons (Fig. 5d), having smaller amplitude, longer half-width and a slower maximum rate of depolarisation and repolarisation (ẟV/ẟt) (Fig. 5e-h). We found no change in AP voltage threshold, rheobase current (Fig. 5i, j) or input resistance (Fig. 5b). The percentage of neurons displaying spontaneous excitatory postsynaptic currents (EPSCs) was comparable at days 50 and 60 (Fig. 5n) as was EPSC frequency and amplitude (Fig. 5l, m). Lack of effect on synaptic transmission may reflect compensation by DLG4, whose expression shows a trend towards an increase in synaptosomes from day 65 *DLG2^-/-^* neurons (Fig. 5o). In summary, developing *DLG2^-/-^* neurons have a reduced ability to fire APs and produce less mature APs.

### Neurogenic programs capture risk variant enrichment in loss-of-function intolerant genes

Having identified neurodevelopmentally expressed pathways enriched for common SZ risk variants and investigated the phenotypic consequences of their dysregulation in *DLG2^-/-^* lines, we sought to test our hypothesis that these pathways capture a significant proportion of the SZ GWAS enrichment seen in LoFi genes^12^. We predicted that LoFi genes would primarily be concentrated in earlier transcriptional programs where the impact of disruption is potentially more severe. LoFi genes were over-represented in all early neurogenic programs but notably depleted in the late set (Fig. 6a). LoFi SZ GWAS association (conditioned on *all^WT+KO^*) was captured by the overlap with early-stable^-/-^ and early-increasing^-/-^, localising the GWAS signal to a fraction of LoFi genes (less than a third) located in specific neurogenic pathways (Fig. 6b).

**Figure 6.**
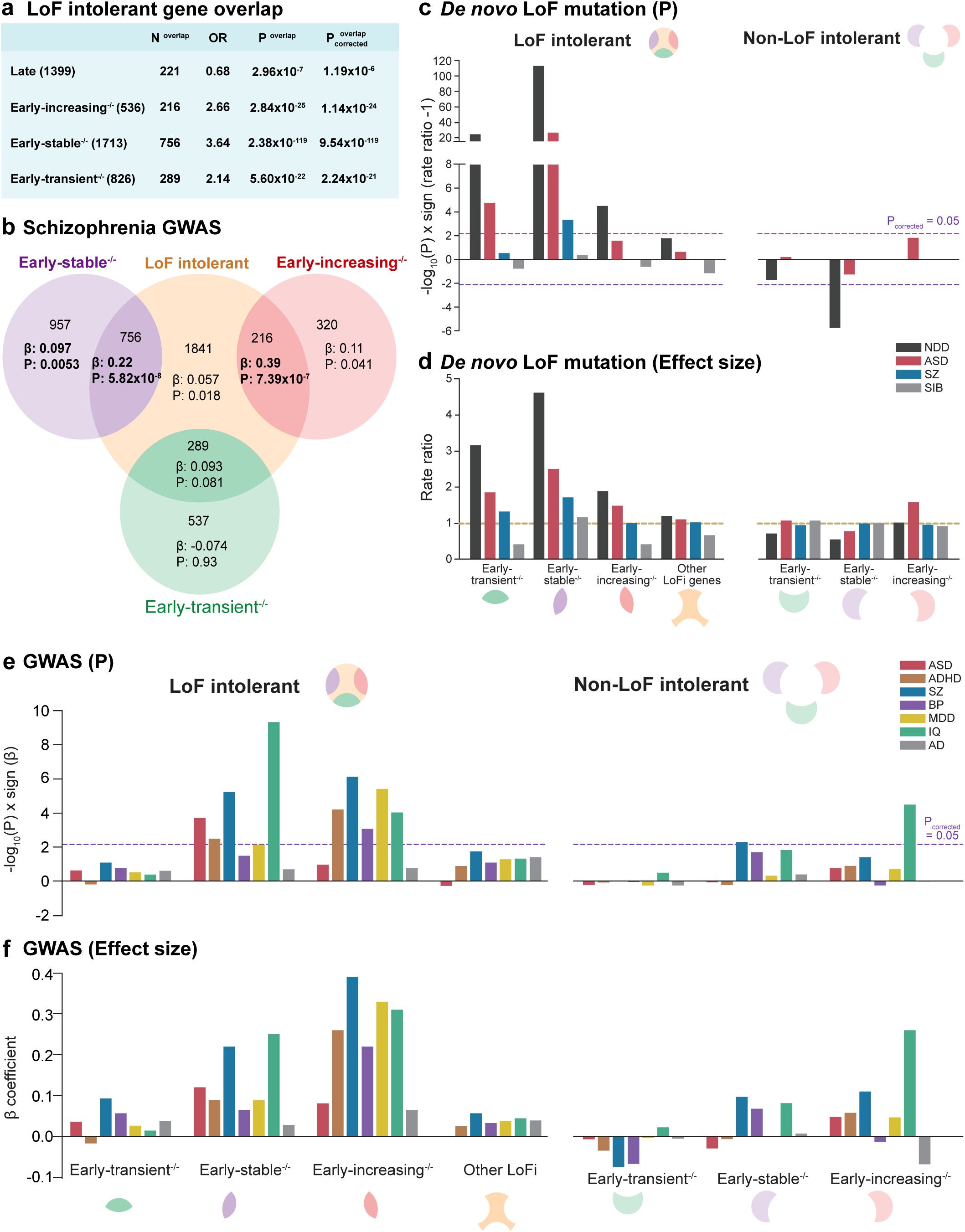
Neuropsychiatric disorder risk variants localise to LoF intolerant genes in early neurogenic transcriptional programs. **a**, Identification of programs enriched for LoFi genes (P^overlap^) relative to all expressed genes. **b**, LoFi genes were partitioned based on their overlap with early neurogenic programs. Each segment of the Venn diagram shows the number of genes in each subset and the regression coefficient (β) and (uncorrected) p-value (P) for SZ common variant enrichment, conditioning on all expressed genes. Bold indicates enrichments surviving correction for 7 tests. **c**, A two-sided Poisson rate ratio test was used to identify programs (partitioned by LoFi gene overlap) enriched for *de novo* LoF mutations from individuals with ID/NDD, ASD and SZ when compared to all other expressed genes. Unaffected siblings (SIB) of ASD cases were analysed as a control. Data points lying above the x axis indicate an increased rate (rate ratio > 1), those below indicate a reduced rate (rate ratio < 1). Dotted lines indicate P_corrected_ = 0.05 following Bonferroni correction for 7 tests (4 LoFi + 3 non-LoFi sets tested for each disorder). **d**, Rate ratios (genes in set versus all other expressed genes) from tests shown in **c**. Dotted line shows rate ratio of 1 (i.e. rate of mutations in set equals that in all other genes). **e**, Programs (partitioned by LoFi gene overlap) were tested for enrichment in common variants contributing to ASD, ADHD, bipolar disorder (BP), major depressive disorder (MDD) and Alzheimer’s disease (AD). Values for SZ (Fig. 6b) included for comparison. Dotted lines indicate P_corrected_ = 0.05 following Bonferroni correction for 7 tests (4 LoFi + 3 non-LoFi sets tested for each disorder). **f**, Effect sizes (β coefficient from MAGMA gene-set enrichment test) for tests shown in **e**.

Under our proposed model, early-transient^-/-^ initiates activation of other early neurogenic programs, thus its dysregulation has the potential to cause more profound developmental deficits. We therefore speculated that – while displaying no evidence for SZ common variant association – LoFi genes in early-transient^-/-^ would be enriched for rare mutations linked to SZ and/or more severe neurodevelopmental disorders. All early neurogenic programs displayed a markedly elevated rate of *de novo* LoF mutations relative to *all^WT+KO^* that was captured by LoFi genes: early-transient^-/-^ for mutations identified in NDD and ASD cases^47^; early-stable^-/-^ for NDD, ASD and SZ^16^; and early-increasing^-/-^ for NDD (Fig. 6c). *De novo* LoF mutations from unaffected siblings of ASD cases^47^ showed no elevation. In all three programs, a clear gradient of effect was evident from NDD (largest elevation in rate) to ASD to SZ, visible only in LoFi genes (Fig. 6d). A modest gradient was also evident for LoFi genes lying outside early neurogenic programs (‘Other LoFi genes’, Fig. 6d), despite *de novo* rates not being robustly elevated here. This suggests the existence of additional biological pathways less central to disease pathophysiology that harbour disease-relevant LoFi genes, although larger samples are likely to be required for their identification.

Given the robust rare variant enrichment across multiple disorders, we investigated whether neurogenic programs are also enriched for common risk variants contributing to disorders other than SZ, analysing a range of conditions with which SZ is known to share heritability: ASD^48^; attention-deficit/hyperactivity disorder (ADHD)^49^; bipolar disorder (BP)^50^; and major depressive disorder (MDD)^51^. Since altered cognitive function is a feature of all these disorders, we also tested programs for enrichment in common variants linked to IQ^52^. Remarkably, all disorders showed evidence for common variant enrichment in one or more early neurogenic program that was again captured by LoFi genes (Fig. 6e, f). In contrast, common variants conferring risk for neurodegenerative disorder Alzheimer’s disease (AD)^53^ were not enriched. Whereas rare variant enrichment was concentrated towards the initial stages of the transcriptional cascade (early-transient^-/-^, early-stable^-/-^), GWAS association was confined to later stages (early-stable^-/-^, early-increasing^-/-^). Conditions with the least prior evidence for an early neurodevelopmental component, BP and MDD, displayed enrichment only in the latest stage (early-increasing^-/-^). Dysregulation of transcriptional programs underlying cortical excitatory neurogenesis thus contributes to a wide spectrum of neuropsychiatric disorders. Furthermore, robust enrichment of early-stable^-/-^ and early-increasing^-/-^ for IQ association (Fig. 6e, f) strongly suggests that perturbation of these neurogenic programs contributes to the emergence of cognitive symptoms in these disorders.

### Divergence between disorders at the level of cellular pathways

For each phenotype-program association identified above, we tested whether GO terms over-represented in that program captured significantly more/less of the association signal than the program as a whole (Methods). As seen for SZ (Fig. 4e, Supplementary Table 6), none of the independent GO term subsets within early-stable^-/-^ or early-increasing^-/-^ showed evidence for depletion of GWAS association (Supplementary Table 7), suggesting that common variants contributing to neuropsychiatric disorders and cognition are distributed across the diverse molecular processes encapsulated by each neurogenic program. However, clear differences in emphasis were evident between disorders: of the two conditions whose association was restricted to early-increasing^-/-^ (Fig. 6e, f), BP showed strong evidence for excess enrichment in *membrane depolarization during action potential* genes – also enriched for SZ GWAS association (Fig. 4e) – but MDD did not (Supplementary Table 7).

Turning to *de novo* rare variant associations, all independent GO term subsets in early-transient^-/-^ (primarily related to transcriptional regulation, Supplementary Table 5) displayed increased enrichment for ASD mutations, whereas excess NDD association was restricted to chromatin binding genes (Supplementary Table 8). Amongst early-stable^-/-^ genes, both ASD and NDD *de novos* were concentrated in transcription factors; in contrast, components of early endosomes were strongly depleted for NDD mutations, as were G-protein-coupled signalling molecules (Guanine nucleotide exchange factors, Rab interactors) – key regulators of endosome function^54, 55^ (Supplementary Table 8). In early-increasing^-/-^, NDD mutations were concentrated in terms related to synaptic transmission and also in *membrane depolarization during action potential* genes (Supplementary Table 8), once again highlighting the importance of this relatively small gene-set.

### Neurogenic programs are active during excitatory corticoneurogenesis in vivo

Genetic analyses (Figs. 4, 6) leave little doubt that early neurogenic programs are highly relevant to cognitive function and the pathogenesis of neuropsychiatric disorders. However, these programs were identified from bulk RNAseq data *in vitro* and it remains to be shown that their constituent genes are actively co-expressed in the appropriate cell-types during cortical excitatory neurogenesis *in vivo*. To address this, we extracted gene expression data for cell-types spanning cortical excitatory neurodevelopment from an existing single-cell RNAseq study of human foetal brain tissue^32^. After normalising the expression for each gene across all cells, we calculated the average expression for each gene in each cell-type/stage of development available: early RG, RG, IPCs, transitioning cells (intermediate between progenitors and neurons), new-born and maturing neurons (Methods). We then plotted the expression of each program (mean and standard error of gene-level averages) in each cell-type/stage and tested for differences in expression between successive types/stages (Fig. 7, Supplementary Table 9). The expression profile seen for each program *in vitro* was recapitulated *in vivo* (Fig. 7). Notably, while other programs in the cascade were significantly upregulated during the transition from progenitors to neurons, early-transient^-/-^ expression was found to be low in early RG, rising in more mature NPCs then declining in neurons. This is consistent with its predicted role in shaping neuronal sub-type identity, which recent evidence indicates is determined by the internal state of NPCs immediately prior to their exit from the cell-cycle^56^.

**Figure 7.**
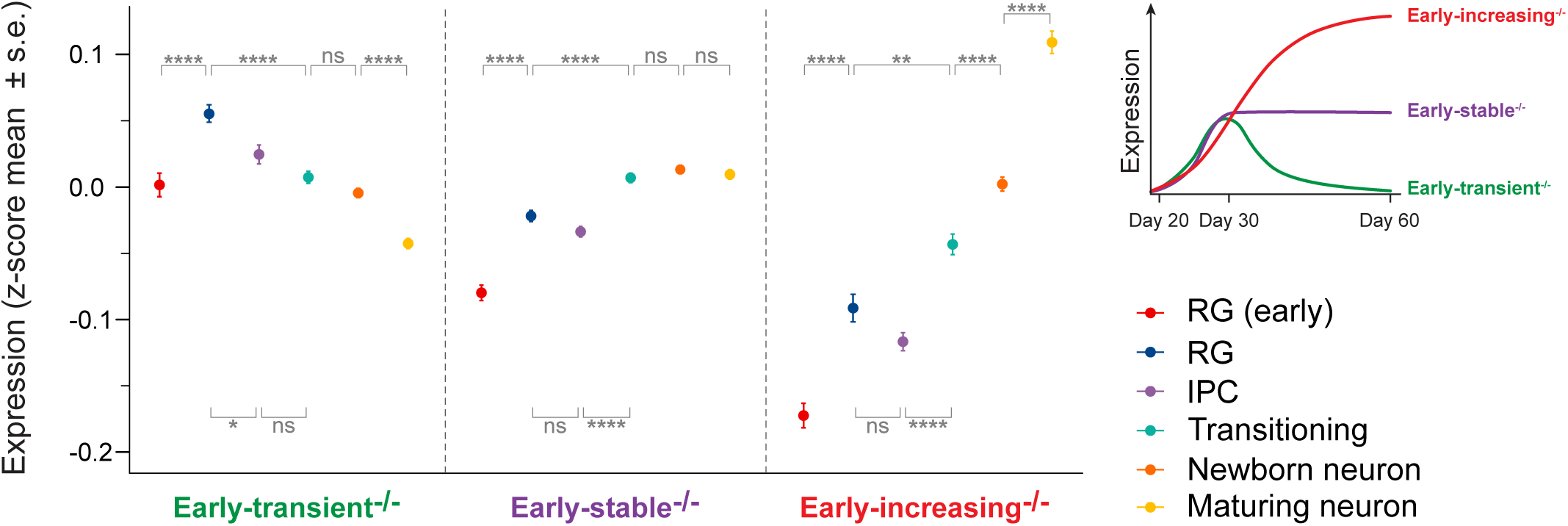
*In vivo* expression of early neurogenic programs in neurodevelopmental cell-types from human foetal cortex. Cells corresponding to distinct neurodevelopmental cell-types (including cell-types at different stages of maturity) were identified and extracted from a previously published single-cell RNA-seq study of human foetal cortex across peak stages of neurogenesis^32^ (Methods). Over 80% of genes belonging to each *in vitro*-defined neurogenic program (top RHS) were present in the *in vivo* data. For each program, the mean and standard error of gene-level expression averages (see main text/Methods) were calculated for each *in vivo* cell-type/developmental stage. The *in vivo* data captures both direct (RG ➝ neuron) and indirect (RG ➝ IPC ➝ neuron) neurogenic pathways. The (deep layer) neurons present in our day 30-60 cultures *in vitro* are predominantly born via the direct neurogenic pathway. For each program, differences between gene-level averages for successive cell-types/stages were evaluated using a two-tailed Student’s t-test and p-values Bonferroni-corrected for the 6 pairwise comparisons. *p<0.05; **p<0.01; ****p<0.0001. All data presented as mean ± SEM.

## Discussion

A complex choreography of cell proliferation, specification, growth, migration and network formation underlies brain development. To date, limited progress has been made pinpointing aspects of this process disrupted in neuropsychiatric disorders. Here we uncover 3 distinct gene expression programs active during early excitatory corticoneurogenesis *in vitro* and *in vivo* (Fig. 7). These programs are highly enriched for variants contributing to a wide spectrum of disorders and cognitive function (Fig. 6). The extent of association across 9 independent genetic datasets is remarkable, each program displaying robust association in multiple studies: 2 for early-transient^-/-^; 7 (including common and rare variation in both ASD and SZ) for early-stable^-/-^; and 6 for early-increasing^-/-^. These programs harbour well-supported risk genes for complex and Mendelian disorders, a number of which are highlighted in Fig. 8a. This convergence of genetic evidence leaves little doubt that these programs play an important aetiological role in a wide range of psychiatric disorders.

**Figure 8.**
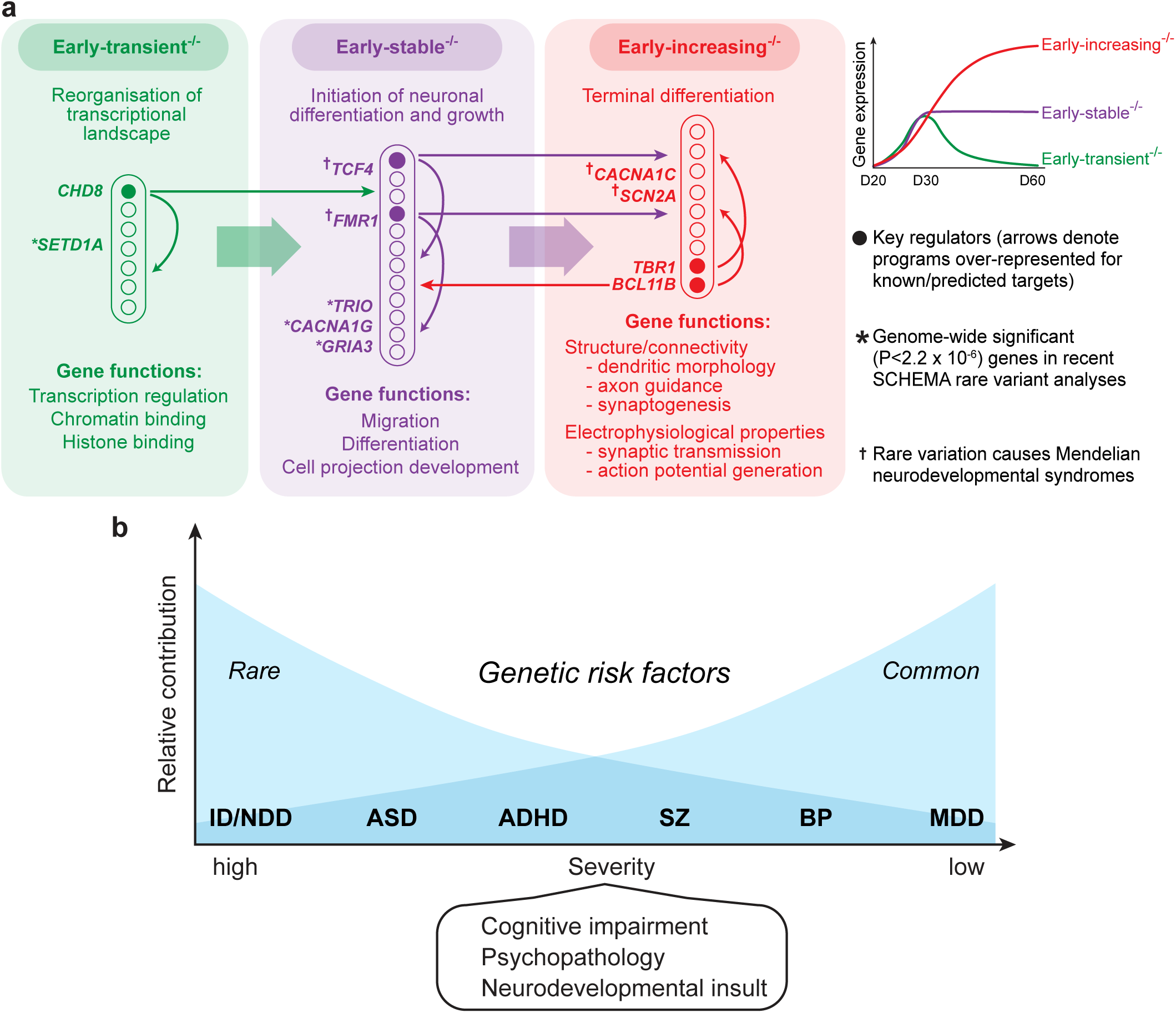
Model of disease pathophysiology in early corticoneurogenesis. **a**, Summary of main GO term enrichments for each transcriptional program (Supplementary Table 5) and identified regulatory interactions between them. Key regulators, genome-wide significant rare coding variants from^79^ and rare mutations causing Mendelian neurodevelopmental syndromes are denoted. **b**, Neurodevelopmental continuum/gradient model. Disorders shown are: intellectual disability/severe neurodevelopmental delay (ID/NDD); autism spectrum disorders (ASD); attention-deficit/hyperactivity disorder (ADHD); schizophrenia (SZ); bipolar disorder (BP); and major depressive disorder (MDD).

Each program has a unique gene expression profile and molecular composition, indicating a distinct functional role during early neurogenesis: based on our findings we propose that they form a transcriptional cascade regulating neuronal growth, migration and differentiation (Fig 8a). Computational analyses of gene/mRNA regulatory interactions implicate known neurodevelopmental disorder risk genes (*CHD8*, *TCF4*, FMRP, *BCL11B* and *TBR1*) as regulators of this cascade and reveal pathways through which they are highly likely to contribute to disease (Fig. 4e). Supporting this model, down-regulation of neurogenic programs in *DLG2*^-/-^ lines is accompanied by deficits that reflect their predicted functions: impaired migration; simplified neuron morphology; immature action potential generation; and delayed expression of cell-type identity (see also Supplementary Discussion). Further experimental work is required to more precisely delineate phenotypes associated with the disruption of individual programs and the risk genes they harbour and to map out regulatory interactions shaping their expression and activity, testing computational predictions. Here we focus on phenotypes expressed by individual new-born excitatory neurons; in future studies it will be important to investigate the persistence of these phenotypes and explore longer-term effects on neuronal circuit formation and function, particularly in light of the predicted role of early-increasing^-/-^ (Fig. 8a).

Cell developmental processes such as growth and migration^57^ arise from the coordinated operation of diverse molecular pathways distributed across multiple organelles. In general, risk variant enrichment in early neurogenic programs did not appear to be restricted to specific functional subsets of genes, suggesting that genetic perturbation of multiple sub-cellular processes is likely to contribute to each disease-relevant cellular phenotype. However, there was evidence that within a given neurogenic program the relative importance of sub-cellular processes varies between disorders. Of the 5 disorders with association in early-increasing^-/-^, SZ, BP and NDD displayed evidence for an excess concentration of genetic risk in a small subset of genes linked to action potential generation, while ADHD and MDD did not. Besides their role in mature neuronal function, mutations in these genes disrupt cortical neuron migration and neurite outgrowth and are implicated in severe neurodevelopmental syndromes (Supplementary Discussion). Within early-stable^-/-^ (predicted to regulate neuronal growth and migration, Fig. 8a) NDD and ASD *de novo* rare variants were strongly concentrated in transcription factors, NDD variants also being highly depleted from genes linked to early endosome function. Amongst other roles, endosomal trafficking regulates the cell-surface expression of receptors and adhesion molecules during migration and axon guidance^55^. It is possible that, even where they impact the same higher level cell developmental process, risk variants for different disorders may display a unique pattern of disruption across underlying molecular pathways.

A clear pattern of enrichment was evident across early neurogenic programs, with rare damaging mutations contributing to more severe disorders concentrated in initial stages (early-transient^-/-^, early-stable^-/-^) of the cascade (Fig. 6d) and common variant association increasing towards later (early-stable^-/-^, early-increasing^-/-^) stages (Fig. 6f). It has been proposed that adult and childhood disorders lie on an aetiological and neurodevelopmental continuum: the more severe the disorder the greater the contribution from rare, damaging mutations and the earlier their developmental impact^58–60^ (Fig. 8b). Our data support this model and ground it in developmental neurobiology, embedding genetic risk for multiple disorders in a common pathophysiological framework.

Genetic risk for all disorders was concentrated in LoFi genes, indicating far wider relevance for these genes than previously appreciated; here we provide the first real insight into their pathophysiological roles. Being under high selective constraint, LoFi genes profoundly impact development through to sexual maturity. It has not been clear whether LoFi genes harbouring pathogenic mutations are distributed across diverse pathways shaping pre-/post-natal growth or are concentrated in specific pathways and/or stages of development. Our analyses reveal that not all neurodevelopmental pathways are enriched for LoFi genes (Fig. 6a), and that the subset of LoFi genes (∼40%) concentrated in early neurogenic programs capture virtually all common and rare variant LoFi association across a wide spectrum of disorders (Fig. 6). While early-transient^-/-^ expression is limited to initial stages of neurogenesis (peaking as radial glia mature, Fig. 7), early-stable^-/-^ and early-increasing^-/-^ continue to be up-regulated during the NPC-neuron transition and persist as neurons mature (Fig. 7), shaping form, function and (potentially) network level organisation at later stages (Fig. 8a).

While it was knockout of *DLG2* that led us to the identification of disease-relevant programs and allowed us to investigate cellular phenotypes associated with their dysregulation, *DLG2* itself has yet to reach the status of a canonical SZ/ASD risk gene. DLG2 is primarily known for its role as a postsynaptic scaffold protein in mature neurons, where it is required for normal formation of NMDA receptor signalling complexes^26^. These complexes are themselves enriched for rare mutations in SZ cases^4, 5, 8–11^, revealing an unexpected link between mature and developmental disease mechanisms. The extent of this link may be far greater than generally appreciated: as noted above, channels involved in action potential generation are also present in early neurogenic programs and are known to impact neuron growth and migration.

Our data reveal that *DLG2* expression is important for cortical excitatory neurodevelopment, but the mechanism by which it operates remains to be uncovered. Based on its known function, and involvement of invertebrate *Dlg* in the developmental *Scrib* signalling module^25^, we propose that DLG2 links cell-surface receptors to signal transduction pathways regulating the activation of neurogenic programs (Supplementary Fig. 9). We hypothesise that stochastic signalling in *DLG2*^-/-^ lines delays and impairs transcriptional activation, disrupting the orchestration of events required for normal development and the specification of neuronal properties. Precise timing is crucial during brain development, where the correct dendritic morphology, axonal length and electrical properties are required for normal circuit formation. Consequently, even transient perturbation of neurogenesis may have a profound impact on fine-grained neuronal wiring, network function and ultimately perception, cognition and behaviour. Clearly much work remains to be done, but we believe that the current findings sketch out a useful neurobiological model upon which future studies into the developmental origins of psychiatric genetic disorders can build.

## Methods

### hESC culture

All hESC lines were maintained at 37°C and 5% CO_2_ in 6 well cell culture plates (Greiner) coated with 1% Matrigel hESC-Qualified Matrix (Corning) prepared in Dulbecco’s Modified Eagle Medium: Nutrient Mixture F-12 (DMEM/F12, Thermo Fisher Scientific). Cells were fed daily with Essential 8 medium (E8, Thermo Fisher Scientific) and passaged at 80% confluency using Versene solution (Thermo Fisher Scientific) for 1.5 minutes at 37°C followed by manual dissociation with a serological pipette. All cells were kept below passage 25 and confirmed as negative for mycoplasma infection.

### DLG2 Knockout hESC line generation

Two guide RNAs targeting exon 22 of the human *DLG2* gene, covering the first PDZ domain, were designed using a web-based tool (crispr.mit.edu) and cloned into two plasmids containing D10A nickase mutant Cas9 with GFP (PX461) or Puromycin resistant gene (PX462)^61^. pSpCas9n(BB)-2A-GFP (PX461) and pSpCas9n(BB)-2A-Puro (PX462) was a gift from Feng Zhang (For PX461, Addgene plasmid#48140; http://n2t.net/addgene:48140; RRID:Addgene_48140; For PX462, Addgene plasmid #48141; http://n2t.net/addgene:48141; RRID:Addgene_48141). H7 hESCs (WiCell) were nucleofected using P4 solution and CB150 programme (Lonza) with 5µg of plasmids, FACS sorted on the following day and plated at a low density (∼70 cells/cm^2^) for clonal isolation. 19 clonal populations were established with 6 WT and 13 mutant lines after targeted sequencing of the exon 22. One WT and two homozygous knockout lines were chosen for study: our WT and KO lines therefore originate from the same H7 parental line and have gone through the same process of nucleofection and FACS sorting together.

### Genetic validation

The gRNA pair had zero predicted off-target nickase sites (Supplementary Fig. 2). Even though we did not use a wild-type Cas9 nuclease (where only a single gRNA is required to create a double-stranded break), we further checked genic predicted off-target sites for each individual gRNA by PCR and Sanger sequencing (GATC & LGC). Out of 30 sites identified, we randomly selected 14 (7 for each gRNA) for validation. No mutations relative to WT were present at any site (Supplementary Table 1). In addition, genotyping on the Illumina PsychArray v1.1 revealed no CNV insertions/deletions in either *DLG2^-/-^* line relative to WT (Supplementary Fig. 5).

### Cortical differentiation

Differentiation to cortical projection neurons (Fig. 1b) was achieved using the dual SMAD inhibition protocol^28^ with modifications (embryoid body to monolayer and replacement of KSR medium with N2B27 medium) suggested by Cambray et al., 2012^29^. Prior to differentiation Versene treatment and mechanical dissociation was used to passage hESCs at approximately 100,000 cells per well into 12 well cell culture plates (Greiner) coated with 1% Matrigel Growth Factor Reduced (GFR) Basement Membrane matrix (Corning) in DMEM/F12, cells were maintained in E8 medium at 37°C and 5% CO_2_ until 90% confluent. At day 0 of the differentiation E8 media was replaced with N2B27-RA neuronal differentiation media consisting of: 2/3 DMEM/F12, 1/3 Neurobasal (Thermo Fisher Scientific), 1x N-2 Supplement (Thermo Fisher Scientific), 1x B27 Supplement minus vitamin A (Thermo Fisher Scientific), 1x Pen Step Glutamine (Thermo Fisher Scientific) and 50 µM 2-Mercaptoethanol (Thermo Fisher Scientific), which was supplemented with 100 nM LDN193189 (Cambridge Biosciences) and 10 µM SB431542 (Stratech Scientific) for the first 10 days only (the neural induction period). At day 10 cells were passaged at a 2:3 ratio into 12 well cell culture plates coated with 15 µg/ml human plasma fibronectin (Merck) in Dulbecco’s phosphate-buffered saline (DPBS, Thermo Fisher Scientific), passage was as previously described with the addition of a 1 hour incubation with 10 µM Y27632 Dihydrochloride (ROCK inhibitor, Stratech Scientific) prior to Versene dissociation. During days 10 to 20 of differentiation cells were maintained in N2B27-RA (without LDN193189 or SB431542 supplementation) and passaged at day 20 in a 1:4 ratio into 24 well cell culture plates (Greiner) sequentially coated with 10 µg/ml poly-d-lysine hydrobromide (PDL, Sigma) and 15 µg/ml laminin (Sigma) in DPBS. Vitamin A was added to the differentiation media at day 26, standard 1x B27 Supplement (Thermo Fisher Scientific) replacing 1x B27 Supplement minus vitamin A, and cells were maintained in the resulting N2B27+RA media for the remainder of the differentiation. Cells maintained to day 40 received no additional passage beyond passage 2 at day 20 while cells kept beyond day 40 received a third passage at day 30, 1:2 onto PDL-laminin as previously described. In all cases cells maintained past day 30 were fed with N2B27+RA supplemented with 2µg/ml laminin once weekly to prevent cell detachment from the culture plates.

### Immunocytochemistry

Cells were fixed in 4% paraformaldehyde (PFA, Sigma) in PBS for 20 minutes at 4°C followed by a 1 hour room temperature incubation in blocking solution of 5% donkey serum (Biosera) in 0.3% Triton-X-100 (Sigma) in PBS (0.3% PBST). Primary antibodies, used at an assay dependent concentration (see ‘Antibody concentration’), were diluted in blocking solution and incubated with cells overnight at 4°C. Following removal of primary antibody solution and 3 PBS washes, cells were incubated in the dark for 2 hours at room temperature with appropriate Alexa Fluor secondary antibodies (Thermo Fisher Scientific) diluted 1:500 with blocking solution. After an additional 2 PBS washes cells were counterstained with DAPI nucleic acid stain (Thermo Fisher Scientific), diluted 1:1000 with PBS, for 5 mins at room temperature and following a final PBS wash, mounted using Dako Fluorescence Mounting Medium (Agilent) and glass coverslips. Imaging was with either the LSM710 confocal microscope (Zeiss) or Cellinsight Cx7 High-Content Screening Platform (Thermo Fisher Scientific) with HCS Studio Cell Analysis software (Thermo Fisher Scientific) used for quantification.

### Western blotting

Total protein was extracted from dissociated cultured cells by incubating in 1x RIPA buffer (New England Biolabs) with added MS-SAFE Protease and Phosphatase Inhibitor (Sigma) for 30 minutes on ice with regular vortexing, concentration was determined using a DC Protein Assay (BioRad) quantified with the CLARIOstar microplate reader (BMG Labtech). Proteins for western blotting were incubated with Bolt LDS sample buffer (Thermo Fisher Scientific) and Bolt Sample Reducing Agent (Thermo Fisher Scientific) for 10 minutes at 70°C before loading into Bolt 4-12% Bis-Tris Plus gels (Thermo Fisher Scientific). Gels were run at 120V for 2-3 hours in Bolt MES SDS Running Buffer (Thermo Fisher Scientific) prior to protein transfer to Amersham Protran nitrocellulose blotting membrane (GE Healthcare) using a Mini Trans-Blot Cell (BioRad) and Bolt Transfer Buffer (Thermo Fisher Scientific) run at 120V for 1 hour 45 minutes. Transfer was confirmed by visualising protein bands with 0.1% Ponceau S (Sigma) in 5% acetic acid (Sigma) followed by repeated H_2_O washes to remove the stain.

Following transfer, membranes were incubated in a blocking solution of 5% milk in TBST, 0.1% TWEEN 20 (Sigma) in TBS (Formedium), for 1 hour at room temperature. Primary antibodies, used at an assay dependent concentration, were diluted with blocking solution prior to incubation with membranes overnight at 4°C. Following 3 TBST washes, membranes were incubated in the dark for 1 hour at room temperature with IRDye secondary antibodies (LI-COR) diluted 1:15000 with blocking solution. After 3 TBS washes staining was visualised using the Odyssey CLx Imaging System (LI-COR).

### Antibody concentration

| Antibody | Company | Identifier Catalog No. | Dilution in ICC | Dilution in WB |
| --- | --- | --- | --- | --- |
| Rabbit polyclonal anti-Calnexin | Abcam | ab22595 | N/A | 1:5000 |
| Rat monoclonal anti-CTIP2 | Abcam | ab18465 | 1:500 | N/A |
| Rabbit polyclonal anti-CTIP2 | Abcam | ab70453 | N/A | 1:1000 |
| Rabbit monoclonal anti-DARPP32 | AbCam | ab40802 | 1:500 | N/A |
| Mouse polyclonal anti-DLX1 | AbCam | ab167575 | 1:100 | N/A |
| Rabbit polyclonal anti-FOXG1 | Abcam | ab18259 | 1:250 | N/A |
| Mouse monoclonal anti-FOXP1 | Abcam | ab32010 | 1:800 | N/A |
| Rabbit polyclonal anti-GABA | Sigma | A2052 | 1:500 | N/A |
| Rabbit polyclonal anti-GAPDH | Abcam | ab9485 | N/A | 1:5000 |
| Goat polyclonal anti-GBX2 | Antibodies.com | A84236 | 1:200 | N/A |
| Mouse monoclonal anti-KI67 | BD Pharmingen | 550609 | 1:150 | 1:1000 |
| Mouse monoclonal anti-MAP2 | Sigma-Aldrich | M1406 | 1:250 | N/A |
| Rabbit polyclonal anti-MAP2 | Merck Millipore | AB5622 | N/A | 1:1000 |
| Rabbit monoclonal anti-NANOG | Cell Signaling Technology | 4903 | 1:400 | N/A |
| Rabbit polyclonal anti-NEUN | Merck Millipore | ABN78 | 1:500 | 1:500 |
| Rabbit monoclonal anti-NKX2.1 | Abcam | ab76013 | 1:1000 | N/A |
| Rabbit monoclonal anti-OCT4 | Cell Signaling Technology | 2840 | 1:400 | N/A |
| Mouse monoclonal anti-OLIG3 | R&D Systems | MAB2456 | 1:500 | N/A |
| Rabbit monoclonal anti-PAX6 | Abcam | ab195045 | 1:300 | N/A |
| Rabbit polyclonal anti-PSD-95 (DLG4) | Abcam | ab18258 | N/A | 1:1000 |
| Mouse monoclonal anti-SATB2 | Abcam | ab51502 | 1:25 | 1:100 |
| Rabbit monoclonal anti-SOX2 | Cell Signaling Technology | 3579 | 1:400 | N/A |
| Goat polyclonal anti-SOX2 | Santa Cruz | sc-17320 | N/A | 1:200 |
| Rabbit polyclonal anti-TBR1 | Abcam | ab31940 | 1:500 | 1:1000 |
| Mouse monoclonal anti-β-Tubulin III (TUJ1) | Sigma-Aldrich | T8660 | 1:1000 | N/A |
| Donkey anti-Rabbit IgG (H+L) secondary antibody, Alexa Fluor 488 | Thermo Fisher Scientific | A-21206 | 1:500 | N/A |
| Donkey anti-Rat IgG (H+L) secondary antibody, Alexa Fluor 488 | Thermo Fisher Scientific | A-21208 | 1:500 | N/A |
| Donkey anti-Mouse IgG (H+L) secondary antibody, Alexa Fluor 594 | Thermo Fisher Scientific | A-21203 | 1:500 | N/A |
| Donkey anti-Rabbit IgG (H+L) secondary antibody, Alexa Fluor 594 | Thermo Fisher Scientific | A-21207 | 1:500 | N/A |
| IRDye 680RD goat anti-Rabbit | Li-Cor | 926-68071 | N/A | 1:15000 |
| IRDye 800CW goat anti-Mouse | Li-Cor | 926-32210 | N/A | 1:15000 |
| IRDye 800CW donkey anti-Goat | Li-Cor | 925-68074 | N/A | 1:15000 |

### Synaptosomal preparation

Synaptic protein was extracted by manually dissociating cultured cells in 1x Syn-PER Reagent (Thermo Fisher Scientific) with added MS-SAFE Protease and Phosphatase Inhibitor (Sigma). Following low speed centrifugation to pellet cell debris (1,200g, 10 min, 4°C) the supernatant was centrifuged at high speed to pellet synaptosomes (15,000g, 20 min, 4°C) which were resuspended in fresh Syn-PER Reagent. Protein concentration was determined using a DC Protein Assay (BioRad) quantified with the CLARIOstar microplate reader (BMG Labtech).

### Peptide affinity purification

PDZ domain containing proteins were enriched from total protein extracts by peptide affinity purification. NMDA receptor subunit 2 C-terminal peptide “SIESDV” was synthesised (Pepceuticals) and fully dissolved in 90% v/v methanol + 1M HEPES pH7 (both Sigma). Dissolved peptide was coupled to Affi-Gel 10 resin (Bio-Rad) that had been washed 3 times in methanol, followed by overnight room temperature incubation on a roller mixer. Unreacted NHS groups were subsequently blocked using 1M Tris pH9 (Sigma) with 2 hours room temp incubation on a roller mixer. The peptide bound resin was then washed 3 times with DOC buffer (1% w/v sodium deoxycholate; 50mM Tris pH9; 1X MS-SAFE Protease and Phosphatase Inhibitor, all Sigma) and stored on ice until required. Total protein was extracted from dissociated cultured cells by incubating in DOC buffer for 1 hour on ice with regular vortexing, cell debris was pelleted by high speed centrifugation (21,300g, 2 hours, 4°C) and the supernatant added to the previously prepared “SIESDV” peptide bound resin. After overnight 4°C incubation on a roller mixer, the resin was washed 5 times with ice cold DOC buffer and the bound protein eluted by 15 minute 70°C incubation in 5% w/v sodium dodecyl sulphate (SDS, Sigma). The eluted protein was reduced with 10 mM TCEP and alkylated using 20 mM Iodoacetamide, trapped and washed on an S-trap micro spin column (ProtiFi, LLC) according to the manufacturer’s instructions and protein digested using trypsin sequence grade (Pierce) at 47°C for 1 hour. Eluted peptides were dried in a vacuum concentrator and resuspended in 0.5% formic acid for MS analysis.

### Mass spectrometry analysis

LC-MS/MS analysis was performed and data was processed and quantified as described previously^62^. Briefly, peptides were analysed by nanoflow LC-MS/MS using an Orbitrap Elite (Thermo Fisher) hybrid mass spectrometer equipped with a nanospray source, coupled to an Ultimate RSLCnano LC System (Dionex). Peptides were desalted on-line using a nano trap column, 75 μm I.D.X 20mm (Thermo Fisher) and then separated using a 130-min gradient from 3 to 40% buffer B (0.5% formic acid in 80% acetonitrile) on an EASY-Spray column, 50 cm × 50 μm ID, PepMap C18, 2 μm particles, 100 Å pore size (Thermo Fisher). The Orbitrap Elite was operated with a cycle of one MS (in the Orbitrap) acquired at a resolution of 60,000 at m/z 400, with the top 20 most abundant multiply charged (2+ and higher) ions in a given chromatographic window subjected to MS/MS fragmentation in the linear ion trap. An FTMS target value of 1e6 and an ion trap MSn target value of 1e4 were used with the lock mass (445.120025) enabled. Maximum FTMS scan accumulation time of 500 ms and maximum ion trap MSn scan accumulation time of 100 ms were used. Dynamic exclusion was enabled with a repeat duration of 45 s with an exclusion list of 500 and an exclusion duration of 30 s. Raw mass spectrometry data were analysed with MaxQuant version 1.6.10.43^63^. Data were searched against a human UniProt sequence database (downloaded December 2019) using the following search parameters: digestion set to Trypsin/P, methionine oxidation and N-terminal protein acetylation as variable modifications, cysteine carbamidomethylation as a fixed modification, match between runs enabled with a match time window of 0.7 min and a 20-min alignment time window, label-free quantification enabled with a minimum ratio count of 2, minimum number of neighbours of 3 and an average number of neighbours of 6. PSM and protein match thresholds were set at 0.1 ppm. A protein FDR of 0.01 and a peptide FDR of 0.01 were used for identification level cut-offs.

### CNV analysis

Following manual dissociation of WT and *DLG2* KO hESC into DPBS, genomic DNA was extracted using the ISOLATE II Genomic DNA kit (Bioline). Following DNA amplification and fragmentation according to the associated Illumina HTS assay protocol samples were hybridized to an Infinium PsychArray v1.1 BeadChip (Illumina). The stained bead chip was imaged using the iScan System (Illuminia) and Genome Studio v2.0 software (Illumina) subsequently used to normalise the raw signal intensity data and perform genotype clustering. Final analysis for Copy Number Variation (CNV) was carried out with PennCNV software^64^.

### RNA sequencing

WT and *DLG2* KO cells were cultured to days 15, 20, 30 and 60 of cortical differentiation as described above (See ‘Cortical differentiation’). Total transcriptome RNA was isolated from triplicate wells for all cell lines at each time point by lysing cells in TRIzol Reagent (Thermo Fisher Scientific) followed by purification with the PureLink RNA Mini Kit (Thermo Fisher Scientific). RNA quality control (QC) was performed with the RNA 6000 Nano kit analysed using the 2100 Bioanalyzer Eukaryote Total RNA Nano assay (Agilent). cDNA libraries for sequencing were produced using the KAPA mRNA HyperPrep Kit for Illumina Platforms (Kapa Biosystems) and indexed with KAPA Single-Indexed Adapter Set A + B (Kapa Biosystems). Library quantification was by Qubit 1x dsDNA HS Assay kit (Thermo Fisher Scientific) and QC by High Sensitivity DNA kit analysed using the 2100 Bioanalyzer High Sensitivity DNA assay (Agilent). Sequencing was performed using the HiSeq4000 Sequencing System (Illumina) with libraries split into 2 equimolar pools, each of which was run over 2 flow cell lanes with 75 base pair paired end reads and 8 base pair index reads.

All samples were modelled after the long-rna-seq-pipeline used by the PsychENCODE Consortium and available at https://www.synapse.org/#!Synapse:syn12026837. Briefly, the fastq files from Illumina HiSeq4000 were assessed for quality by using FastQC tool (v0.11.8)^65^ and trimmed for adapter sequence and low base call quality (Phred score < 30 at ends) with cutadapt (v2.3)^66^. The mapping of the trimmed reads was done using STAR (v2.7.0e)^67^ and the BAM files were produced in both genomic and transcriptomic coordinates and sorted using samtools (v1.9)^68^. The aligned and sorted BAM files were further assessed for quality using Picard tools (v2.20.2)^69^. This revealed a high level of duplicate reads in day 30 KO2 samples (∼72% compared to an average of 23% for other samples). These samples were removed prior to further analyses, which were thus performed on KO1 and WT samples for this timepoint. GRCh38.p13 was used as the reference genome and the comprehensive gene annotations on the primary assembly from Gencode (release 32) used as gene annotation. Gene and transcript-level quantifications were calculated using RSEM (v1.3.1)^70^. Both STAR and RSEM executions were performed using the psychENCODE parameters.

RSEM gene and isoform level estimated counts were imported using the tximport package (v1.12.3)^71^. Protein coding genes expressed (cpm>=1) in at least 1/3 of the samples were taken forward for differential analyses of genes, transcripts and exons. Differential gene expression analysis was performed using the DESeq2 package (v1.24.0)^72^ and differentially expressed genes were considered significant if their p value after Bonferroni correction was < 0.05. Differential exon usage was analysed using the DEXSeq pipeline^73^. Briefly, the GENCODE annotation .gtf file was translated into a .gff file with collapsed exon counting bins by using the dexseq_prepare_annotation.py script. Mapped reads overlapping each of the exon counting bins were then counted using the python_count.py script and the HTSEQ software (0.11.2)^74^. Finally, differential exon usage was evaluated using DEXSeq (v1.30)^73^ and significant differences identified using an FDR threshold of 0.05. All the differential analyses were performed by using R (v3.6.1).

When analysing differential gene expression in *DLG2^-/-^* relative to WT, samples from KO1 and KO2 lines were combined i.e. for each timepoint a single differential gene expression analysis was performed, comparing expression in KO1 & KO2 samples against wild-type. To assess the impact of sample dropout at day 30, we investigated the similarity in gene expression between lines by clustering all KO1, KO2 and WT samples (Supplementary Fig. 3a). At all 4 timepoints, all replicates from KO1 and KO2 cluster together: while KO2 samples from day 30 are not of sufficient quality to reliably inform further analyses, they are clearly similar to KO1 day 30 samples. We also performed differential expression analyses separately for each line (i.e. KO1 v WT and KO2 v WT) at all other timepoints. The overlap in expressed genes accounted for >98% of the genes expressed in each line and gene expression fold change was highly correlated between KO1 v WT and KO2 v WT (Spearman’s ⍴^day 15^ = 0.67, ⍴ ^day 20^ = 0.95, ⍴ ^day 60^ = 0.75). Over-representation odds ratios for GO terms also remain well correlated for significantly up-regulated (⍴^day 15^ = 0.70, ⍴ ^day 20^ = 0.92, ⍴ ^day 60^ = 0.67) and down-regulated regulated (⍴^day 15^ = 0.55, ⍴^day 20^ = 0.95, ⍴^day 60^ = 0.56) genes. We noted that agreement between lines was greatest for day 20, which also lies close to the onset of neurogenesis and displays a high level of differential expression between KO and WT lines, comparable to that for day 30 (Figure 1g). Further indicating a limited impact for sample dropout, phenotypes predicted by GO term analysis of differential expression at day 30 (deficits in neuron migration, morphology and action potential generation) were experimentally validated (Fig. 3 & 5); and all early neurogenic transcriptional programs identified in these data were shown to possess an identical profile of expression across human neurodevelopmental cell-types *in vivo* (Fig. 7).

### Transcriptional programs

Genes were partitioned based upon their WT expression profiles as follows. Differentially expressed genes (Bonferroni P < 0.05) were first identified between pairs of timepoints (analysing WT data only): genes up-regulated in day 30 relative to day 20 (20-30_up_^WT^); genes up-regulated in day 60 relative to day 30 (30-60_up_^WT^); and genes up-regulated in day 60 relative to day 20 (20-60_up_^WT^). Early-transient, early-stable, early-increasing and late programs were then defined based upon the intersections of these gene-sets as shown in Fig. 4b.

### Human foetal cortex single-cell RNA sequencing data

Single-cell RNA-Seq gene expression data from Nowakowski et al., 2017^32^ were downloaded from <u>bit.ly/cortexSingleCell</u>. Cells corresponding to distinct neurodevelopmental cell-types (including cell-types at different stages of maturity) were identified and extracted, collating all cells from the corresponding *in vivo* cell clusters^32^ as follows:

#### Progenitors

RG (early): “RG-early”

RG: “RG-div1”, “RG-div2”, “oRG”, “tRG”, “vRG” IPC: “IPC-div1”, “IPC-div2”

*Transitioning*: “IPC-nEN1”, “IPC-nEN2”, “IPC-nEN3”

#### Cortical excitatory neurons

Newborn: “nEN-early1”, “nEN-early2”, “nEN-late”

Maturing: “EN-PFC1”,“EN-PFC2”,“EN-PFC3”,“EN-V1-1”,“EN-V1-2”, “EN-V1-3”

Cells with less than 5% of all protein-coding genes expressed (TPM>0) and genes expressed in less than 5% of cells were filtered out. The remaining dataset consisted of 2318 cells and 9239 protein-coding genes. Gene expression counts (TPM) were z-score normalised for each gene across all cells, then the average normalised expression score for each gene was calculated for each of the above cell-types. Over 80% of genes for each *in vitro* program (early-transient^-/-^, early-stable^-/-^ and early-increasing^-/-^) were present in the *in vivo* data; all of these genes passed our filtering criteria. Taking the set of genes corresponding to each *in vitro* program, we calculated the mean and standard error of their gene-level averages in each *in vivo* cell-type (Fig. 7). For each program, the difference between successive neurodevelopmental cell-types/stages was calculated using a two-tailed Student’s t-test and p-values Bonferroni-corrected for the 6 comparisons made: RG (early) v RG; RG v IPC; RG v Transitioning; IPC v Transitioning; Transitioning v Newborn neurons; and Newborn v Maturing neurons.

### Gene set construction

#### GO

The Gene Ontology (GO) ontology tree was downloaded from OBO: http://purl.obolibrary.org/obo/go/go-basic.obo

Ontology trees were constructed separately for Molecular Function, Biological Process and Cellular Component using ‘is_a’ and ‘part_of’ relationships. GO annotations were downloaded from NCBI:

ftp://ftp.ncbi.nlm.nih.gov/gene/DATA/gene2go.gz

Annotations containing the negative qualifier NOT were removed, as were all annotations with evidence codes IEA, NAS and RCA. Annotations were further restricted to protein-coding genes. Genes corresponding to each annotation term were then annotated with all parents of that term, identified using the appropriate ontology tree. Finally, terms containing between 20 and 2000 genes were extracted for analysis.

#### Regulator targets

##### Predicted *TBR1* and *BCL11B* targets^42^

Transcription factor-target gene interactions identified by elastic net regression were downloaded from the PsychEncode resource website (http://resource.psychencode.org/#Derived) and predicted targets for *TBR1* and *BCL11B* extracted (interaction file: INT-11_ElasticNet_Filtered_Cutoff_0.1_GRN_1.csv). Gene symbols were mapped to NCBI/Entrez ids using data from the NCBI gene_info file.

##### TCF4 targets^40^

identifiers were updated using the gene_history file from NCBI.

##### FMRP targets^41^

NCBI/Entrez mouse gene identifiers were updated using the gene_history file from NCBI. Genes were then mapped from mouse to human using Homologene, restricting to protein-coding genes with a 1-1 mapping.

##### Direct *CHD8* mid-foetal promoter targets^37^

symbols (NCBI gene_info file) and locations (NCBI Build37.3) were used to map genes to NCBI/Entrez gene ids. Note: using Sugathan et al., 2014^36^ rather than Cotney et al., 2015^37^ to define direct CHD8 targets did not alter the observed pattern of overlap with transcriptional programs (Fig. 4)(data not shown).

##### Indirect *CHD8* targets^36^

Ensembl ids were updated (Ensembl stable_id_event file) then mapped to current NCBI/Entrez ids using Ensembl and NCBI id cross-reference files. Taking genes with altered expression on *CHD8* shRNA knockdown, we removed those identified as direct *CHD8* targets in NPCs^36^ or as *CHD8* mid-foetal promoter targets^37^. Using a stricter definition of genes with altered expression on *CHD8* shRNA knockdown than that taken by Sugathan et al., 2014^36^ – genes with a Bonferroni corrected differential expression P < 0.05, rather than genes with a nominal P < 0.05 – did not alter the pattern of overlap with transcriptional programs (data not shown).

### Functional over-representation test (gene set overlap)

The degree of overlap between pairs of gene sets was evaluated using Fisher’s Exact test, where the background set consisted of all genes expressed in either WT or *DLG2*^-/-^ lines (*all^WT+KO^*). This was used for GO terms; the analysis of regulator targets (Fig. 4e); and the overlap between LoFi genes and transcriptional programs (Fig. 6a). In order to identify a semi-independent subset of over-represented annotations from the output of GO term tests (Fig. 4f, Supplementary Table 4, 5), we used an iterative refinement procedure. Briefly, we selected the gene set with the largest enrichment odds ratio; removed all genes in this set from all other over-represented annotations; re-tested these reduced gene-sets for over-representation in 30_down_^-/-^ genes; then discarded gene-sets with P ≥ 0.05 (after Bonferroni-correction for the number of sets tested in that iteration). This process was repeated (with gene-sets being cumulatively depleted of genes at each iteration) until there were no remaining sets with a corrected P < 0.05.

### Common variant association

All common variant gene-set enrichment analyses were performed using the competitive gene-set enrichment test implemented in MAGMA version 1.07, conditioning on *all^WT+KO^* using the *condition-residualize* function. To test whether GO terms (Fig 4f, Supplementary Table 6) or regulator targets (Fig. 4e) enriched in a specific program captured more or less of the SZ association in these programs than expected, a two-sided enrichment test was performed on term/target genes within the program, conditioning on *all^WT+KO^* and on all genes in the program. All other GWAS enrichment tests were one-sided. To test whether common variant enrichment differed between two gene-sets, we took the regression coefficient β and its standard error SE(β) for each gene-set from the MAGMA output file and compared z = d/SE(d) to a standard normal distribution, where d = β_1_ – β_2_ and SE(d) = √[SE(β_1_)^2^ + SE(β_2_)^2^]. Gene-level association statistics for schizophrenia were taken from Pardiñas et al., 2018^12^; those for ADHD^49^, bipolar disorder^50^ and Alzheimer’s disease^53^ were calculated using the MAGMA multi model, with a fixed 20,000 permutations for each gene. Prior to analysis, SNPs with MAF < 0.01 or INFO score < 0.6 were removed from the bipolar GWAS, bringing it into line with the other datasets.

### Rare variant association

The *de novo* LoF mutations for SZ analysed here are described in Rees et al., 2020^16^. *De novo* LoF mutations for NDD, ASD and unaffected siblings of individuals with ASD were taken from Satterstrom et al., 2020^47^: these were re-annotated using VEP^75^ and mutations mapping to > 2 genes (once readthrough annotations had been discarded) were removed from the analysis. A two-sided Poisson rate ratio test was used to evaluate whether the enrichment of *de novo* LoF mutations in specific gene-sets was significantly greater than that observed for all other expressed genes (using *all^WT+KO^*). The expected rate of *de novo* LoF mutations in a set of genes was estimated using individual gene mutation rates^76^.

### Migration assay

Cells were cultured and differentiated to cortical projection neurons as previously described. Neuronal migration was measured during a 70-hour period from day 40 by transferring cell culture plates to the IncuCyte Live Cell Analysis System (Sartorius). Cells were maintained at 37°C and 5% CO_2_ with 20X magnification phase contrast images taken of whole wells in every 2 hours for the analysis period. The StackReg plugin^77^ for ImageJ was used to fully align the resulting stacks of time lapse-images after which the cartesian coordinates of individual neuronal soma were recorded over the course of the experiment, enabling the distance and speed of neuronal migration to be calculated. Data sets (Fig. 3f, g) were analysed by unpaired two-tailed Student’s t-test.

### Morphology analysis

Cells were differentiated to cortical projection neurons essentially as described and neuronal morphology assessed at days 30 and 70. To generate low density cultures for analysis, cells were passaged at either day 25 or 50 using 15-minute Accutase solution (Sigma) dissociation followed by plating at 100,000 cells per well on 24 well culture plates. 72 hours prior to morphology assessment cells were transfected with 500ng pmaxGFP (Lonza) per well using Lipofectaime 3000 Reagent (Thermo Fisher Scientific) and Opti-MEM Reduced Serum Media (Thermo Fisher Scientific) for the preparation of DNA-lipid complexes. At days 30 or 70, cells were fixed in 4% paraformaldehyde (PFA, Sigma) in PBS for 20 minutes at 4°C before mounting with Dako Fluorescence Mounting Medium (Agilent) and glass coverslips. Random fields were imaged using a DMI6000B Inverted microscope (Leica) and the morphology of GFP expressing cells with a clear neuronal phenotype quantified using the Neurolucida 360 (MBF Bioscience) neuron tracing and analysis software package. Data sets (Fig. 3a-d) were analysed by two-way ANOVA with post hoc comparisons using Bonferroni correction, comparing to WT controls.

### Electrophysiology

Whole cell patch clamp electrophysiology was performed on cells cultured on 13mm round coverslips and the most morphologically mature neurons were patched in each culture; hence the most comparable subpopulation of cells from each genotype was compared. On day 20 of hESC differentiation, 250,000 human neural precursor cells from WT and KO hESCs were dissociated and plated on each PDL-coated coverslip in 30µl diluted (20x) matrigel (Corning) together with 20,000 rat primary glial cells. Postnatal day 7-10 old Sprague-Dawley rats (Charles River) bred in-house were sacrificed via cervical dislocation and cortex was quickly dissected. Tissues were dissociated using 2mg/ml papain and plated in DMEM supplemented with 10% Foetal bovine serum and 1% penicillin/streptomycin/Amphotericin B and 1x Glutamax (all Thermo Fisher Scientific). Microglia and oligodendrocyte precursor cells were removed by shaking at 500 rpm for 24 hours at 37°C. All animal procedures were performed in accordance with Cardiff University’s animal care committee’s regulations and the European Directive 2010/63/EU on the protection of animals used for scientific purposes. Plated cells were fed with BrainPhys medium (Stem cell Technologies) supplemented with 1x B27 (Thermo Fisher Scientific), 10ng/ml BDNF (Cambridge Bioscience) and 200µM ascorbic acid (Sigma). To stop the proliferation of cells, 1x CultureOne (Thermo Fisher Scientific) was supplemented from day 21. For postsynaptic current experiment, coverslips were transferred to a recording chamber (RC-26G, Warner Instruments) and perfused with HEPES Buffered Saline (HBS) (119 mM NaCl; 5 mM KCl; 25 mM HEPES; 33 mM glucose; 2mM CaCl_2_; 2mM MgCl_2_; 1µM glycine; 100µM picrotoxin; pH 7.4), at a flow rate of 2-3 ml per minute. Recordings were made using pipettes pulled from borosilicate glass capillaries (1.5 mm OD, 0.86 mm ID, Harvard Apparatus), and experiments were performed at room temperature (∼20 °C). mEPSC recordings were made using recording electrodes filled with a Cs-based intracellular filling solution (130 mM CsMeSO_4_; 8 mM NaCl; 4 mM Mg-ATP; 0.3 mM Na-GTP, 0.5 mM EGTA; 10 mM HEPES; 6 mM QX-314; with pH 7.3 and osmolarity ∼295 mOsm). Cells were voltage clamped at −60 mV using a Multiclamp 700B amplifier (Axon Instruments). Continuous current acquisition, series resistance and input resistance were monitored and analysed online and offline using the WinLTP software^78^ (http://www.winltp.com). Only cells with series resistance <25 MΩ with a change in series resistance <10% from the start were included in this study. Data were analysed by importing Axon Binary Files into Clampfit (version 10.6; Molecular Devices). A threshold function of >12 pA was used to identify mEPSC events, which were then subject to manual confirmation. Results were outputted to SigmaPlot (version 12.5, Systat Software), where analysis of peak amplitude and frequency of events was performed. The current clamp was used to record resting membrane potential (RMP) and action potentials (AP). Data were sampled at 20kHz with a 3 kHz Bessel filter with MultiClamp 700B amplifier. Coverslips were transferred into the recording chamber maintained at RT (20-21°C) on the stage of an Olympus BX61W (Olympus) differential interference contrast (DIC) microscope and perfused at 2.5ml/min with the external solution composed of 135mM NaCl, 3mM KCl, 1.2mM MgCl_2_, 1.25mM CaCl_2_, 15mM D-glucose, 5mM HEPES (all from Sigma), and pH was titrated to 7.4 by NaOH. The internal solution used to fill the patch pipettes was composed of 117mM KCl, 10mM NaCl, 11mM HEPES, 2mM Na_2_-ATP, 2mM Na-GTP, 1.2mM Na_2_-phosphocreatine, 2mM MgCl_2_, 1mM CaCl_2_ and 11mM EGTA (all from Sigma), and pH was titrated to 7.3 by NaOH. The resistance of a patch pipette was 3–9 MΩ and the series resistance component was fully compensated using the bridge balance function of the instrument. The RMP of cells was recorded immediately after breaking into the cells in gap free mode. A systematic current injection protocol (duration, 1 s; increment, 20 pA; from - 60pA to 120pA) was applied to the neurons held at −60mV to evoke APs. Input resistance (R_in_) was calculated by R_in_=(V_i_-V_m_)/I, where V_i_ is the potential recorded from -10pA current step. The AP properties are measured by the first over shooting AP. Further analysis for action potential characterization was carried out by Clampfit 10.7 software (Molecular Devices).

### Statistical analysis and data presentation

Unless specifically stated in each methodology section, GraphPad Prism (version 8.3.0) was used to test the statistical significance of the data and to produce the graphs. Stars above bars in each graph represents Bonferroni-corrected post hoc tests, *P<0.05; **P<0.01; ***P<0.001; ****P<0.0001 vs. WT control. All phenotypic validation results were from a minimum of two independent differentiations unless otherwise stated, within a given differentiation triplicate samples were used per cell line at each time point investigated. All data presented as mean ± SEM.

### Bonferroni test correction

GO term analyses were corrected for the ∼4,200 terms tested.

Fig. 2a – corrected for 8 tests (up & down regulated x 4 timepoints)

Fig. 4a – corrected for 6 tests (up & down regulated x 3 pairs of timepoints) Fig. 4c – corrected for 7 tests (7 gene expression programs)

Fig. 4d (P^overlap^) – corrected for 21 tests (7 regulator target sets x 3 programs)

Fig. 4d (P^GWAS^) – corrected for 10 tests (10 over-represented sets taken forward for genetic analysis)

Fig. 4e – corrected for 16 tests (16 semi-independent over-represented GO terms) Fig. 6a – corrected for 4 tests (4 programs)

Fig. 6b – uncorrected (secondary, exploratory analysis: results in **bold** survive correction for 7 tests)

Fig. 6c – each disorder (& SIB controls) corrected for 7 tests (4 LoFi + 3 non-LoFi gene-sets) Fig. 6e – each disorder corrected for 7 tests (4 LoFi + 3 non-LoFi gene-sets)

Fig. 7 – each program (early-transient^-/-^, early-stable^-/-^, early-increasing^-/-^) corrected for 6 tests Supplementary Table 6 – corrected for 13 tests (13 semi-independent over-represented GO terms)

Supplementary Table 7 – each disorder corrected for number of GO terms analysed (13 early-stable^-/-^, 16 early-increasing^-/-^)

Supplementary Table 8 – each disorder corrected for number of GO terms analysed (4 early-transient^-/-^, 13 early-stable^-/-^, 16 early-increasing^-/-^)

Supplementary Table 9 – each program (early-transient^-/-^, early-stable^-/-^, early-increasing^-/-^) corrected for 6 tests

### Data usage acknowledgements

We thank the International Genomics of Alzheimer’s Project (IGAP) for providing summary results data for AD common variant analysis. The investigators within IGAP contributed to the design and implementation of IGAP and/or provided data but did not participate in analysis or writing of this report. IGAP was made possible by the generous participation of the control subjects, the patients, and their families. The i–Select chips was funded by the French National Foundation on Alzheimer’s disease and related disorders. EADI was supported by the LABEX (laboratory of excellence program investment for the future) DISTALZ grant, Inserm, Institut Pasteur de Lille, Université de Lille 2 and the Lille University Hospital. GERAD was supported by the Medical Research Council (Grant n° 503480), Alzheimer’s Research UK (Grant n° 503176), the Wellcome Trust (Grant n° 082604/2/07/Z) and German Federal Ministry of Education and Research (BMBF): Competence Network Dementia (CND) grant n° 01GI0102, 01GI0711, 01GI0420. CHARGE was partly supported by the NIH/NIA grant R01 AG033193 and the NIA AG081220 and AGES contract N01–AG–12100, the NHLBI grant R01 HL105756, the Icelandic Heart Association, and the Erasmus Medical Center and Erasmus University. ADGC was supported by the NIH/NIA grants: U01 AG032984, U24 AG021886, U01 AG016976, and the Alzheimer’s Association grant ADGC–10–19672

We thank the research participants and employees of 23andMe for the sharing of summary statistics for MDD common variant analysis.

## Supporting information

Supplementary Discussion

## Acknowledgments

This work was supported by Wellcome Trust Strategic Award (100202/Z/12/Z), MRC programme grant (G08005009), MRC Centre grant (MR/L010305/1), Waterloo Foundation ‘Changing Minds’ programme and start-up funding from the Neuroscience and Mental Health Research Institute, Cardiff University. We acknowledge excellent technical support for RNA sequencing from Joanne Morgan (MRC Centre) and MS analysis from Lydia Kiesel (University of Sheffield) and assistance in morphology tracing from Sophie Pocklington. We appreciate excellent general lab support from Emma Dalton, Trudy Workman and Olena Petter. We thank Prof. Meng Li for her advice and Dr. Claudia Tamburini for technical support in the initial stages of the project and Profs. Yves Barde and Lesley Jones for helpful comments on the manuscript and Emily Adair for providing rat primary glial cells. For data usage acknowledgements, see Methods.

## Author contributions

Conceptualization, AP, ES

Methodology, AE, DB, AP, ES

Software/Data curation, DDA, AP

Formal analysis/Investigation, BS, DDA, MC, TS, ER, YZ, GC, SL, AFP, DW, AP, ES

Writing – Original Draft, BS, DDA, MC, YZ, DW, AP, ES Writing –

Review & Editing, BS, DDA, MC, ER, MOD, MO, AE, DB, DW, AP, ES

Visualization, BS, DDA, MC, YZ, DW, AP, ES

Supervision, AP, ES

Funding acquisition, AH, MOD, MO, AP, ES

## Correspondence and requests for materials

should be addressed to E.S or A.J.P.

## Data availability

RNAseq data generated by this study have been deposited in the European Nucleotide Archive with the accession number PRJEB35773. Additional data that support the findings of this study but are not included in the manuscript, figures and supplementary information are available from the corresponding author upon request.

## Code availability

All publicly available software utilised are noted in Methods. The custom R script used to perform GO term over-representation and refinement analyses is available from GitHub (https://github.com/ajp-cdf/Gene-set-over-representation-refinement).

## Supplemental Information

**Supplementary Figure 1.**
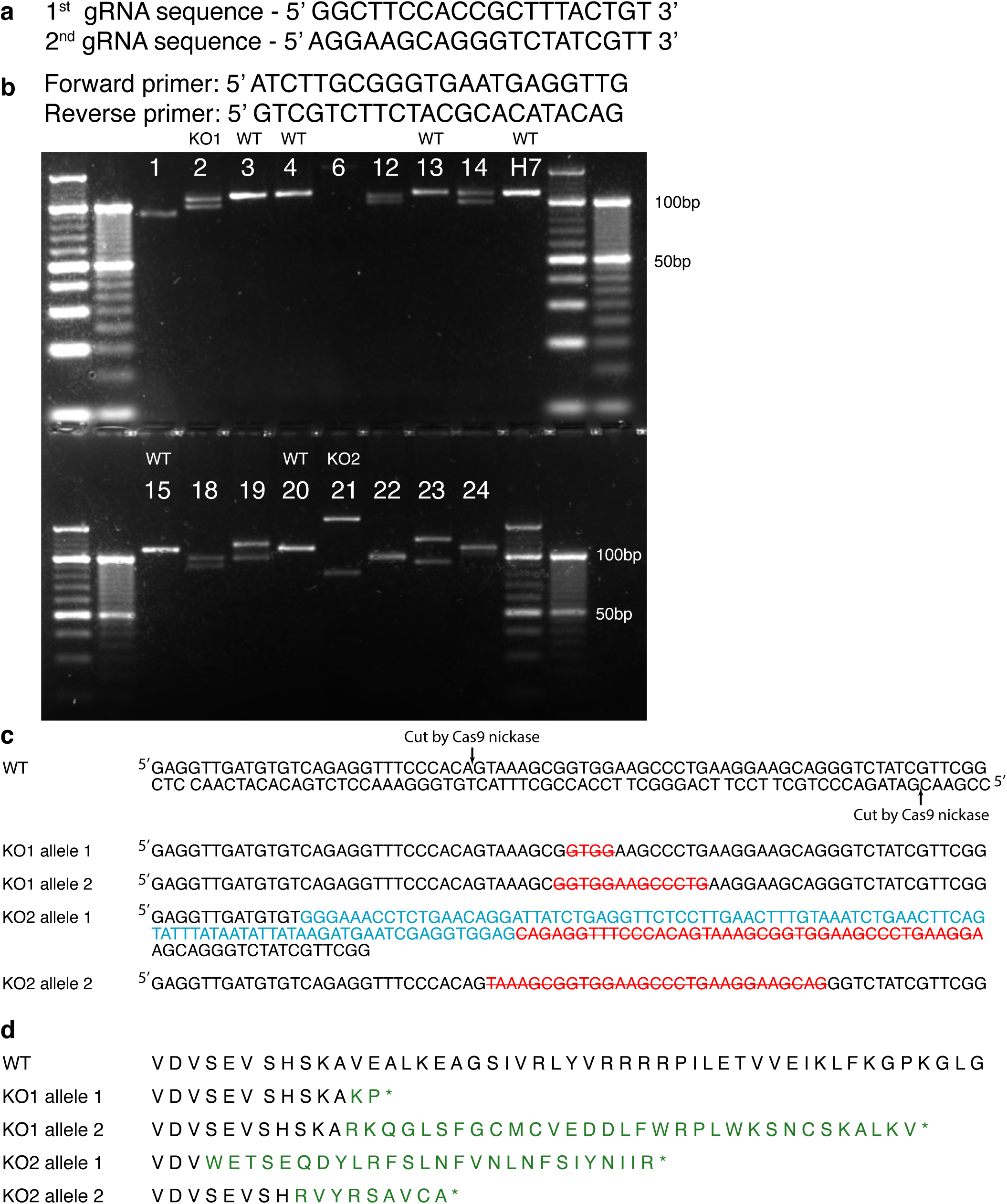
Generation of *DLG2^-/-^* human embryonic stem cell lines. **a**, Two guide RNA sequences used to target exon 22 of *DLG2* which is translated into part of the 1st PDZ domain. **b**, A primer set used for screening successful indels on targeted DNA. PCR and gel electrophresis shows several hESC clones with indels. Parental H7 lines shows a single 109bp band whereas several clones, such as 2 and 21, show two different sized bands suggesting generation of indels on the targted exon. **c**, PCR amplicons were sequenced and clones no.2 and 21 found to be homozygous knockouts (named KO1 and KO2). Red letters indicate deleted bases while blue indicate inserted DNA. **d**, Indels in KOs generated premature stop codons on the targeted exon. Green indicates different amino acid sequences from WT.

**Supplementary Figure 2.**
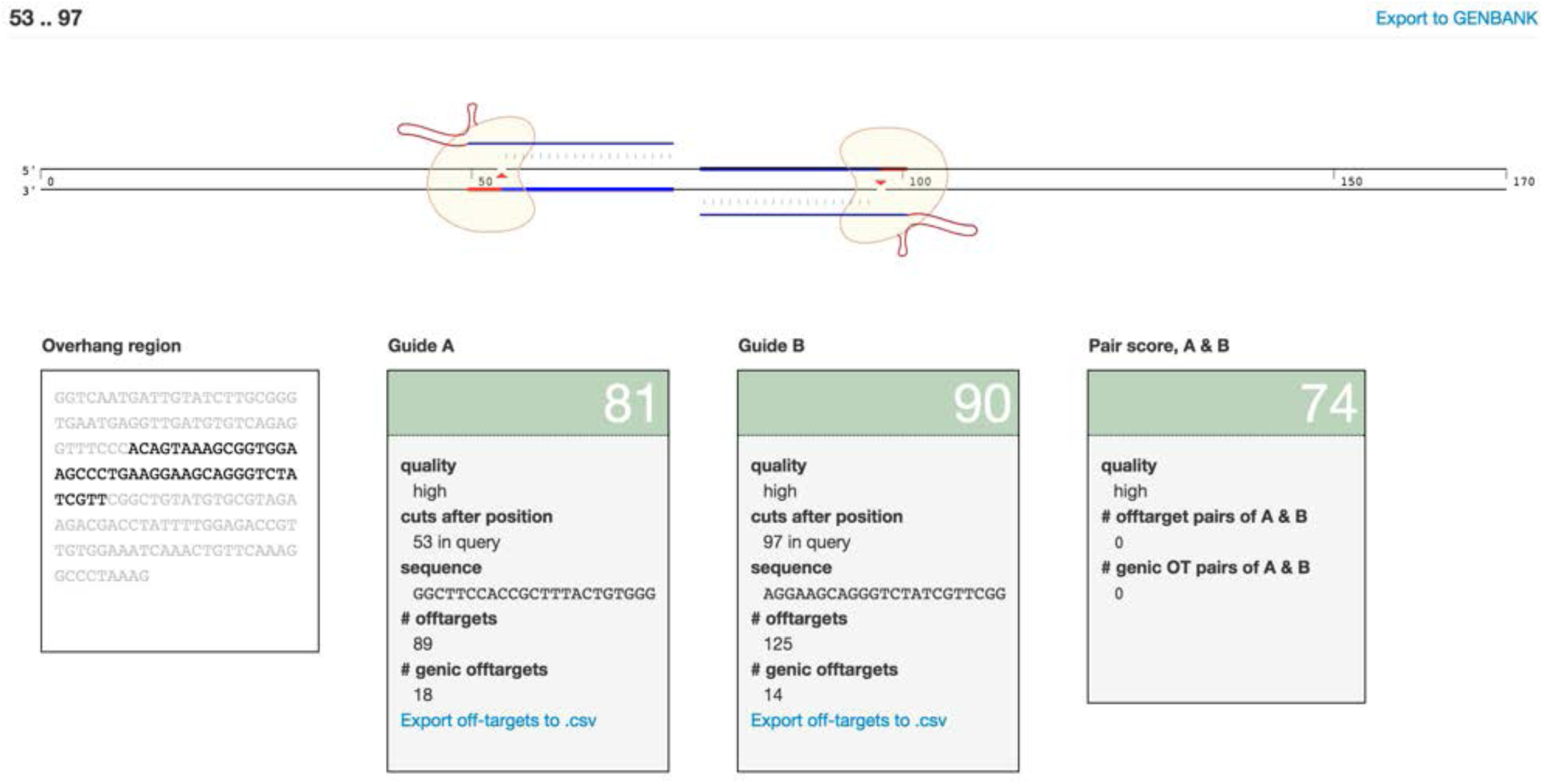
Quality of two gRNAs used in the study. Seperately the gRNAs had prediced off-targets of 89 and 125 loci each; however, the predicted off-targets when used as a pair was none. Captured from crispr.mit.edu.

**Supplementary Figure 3.**
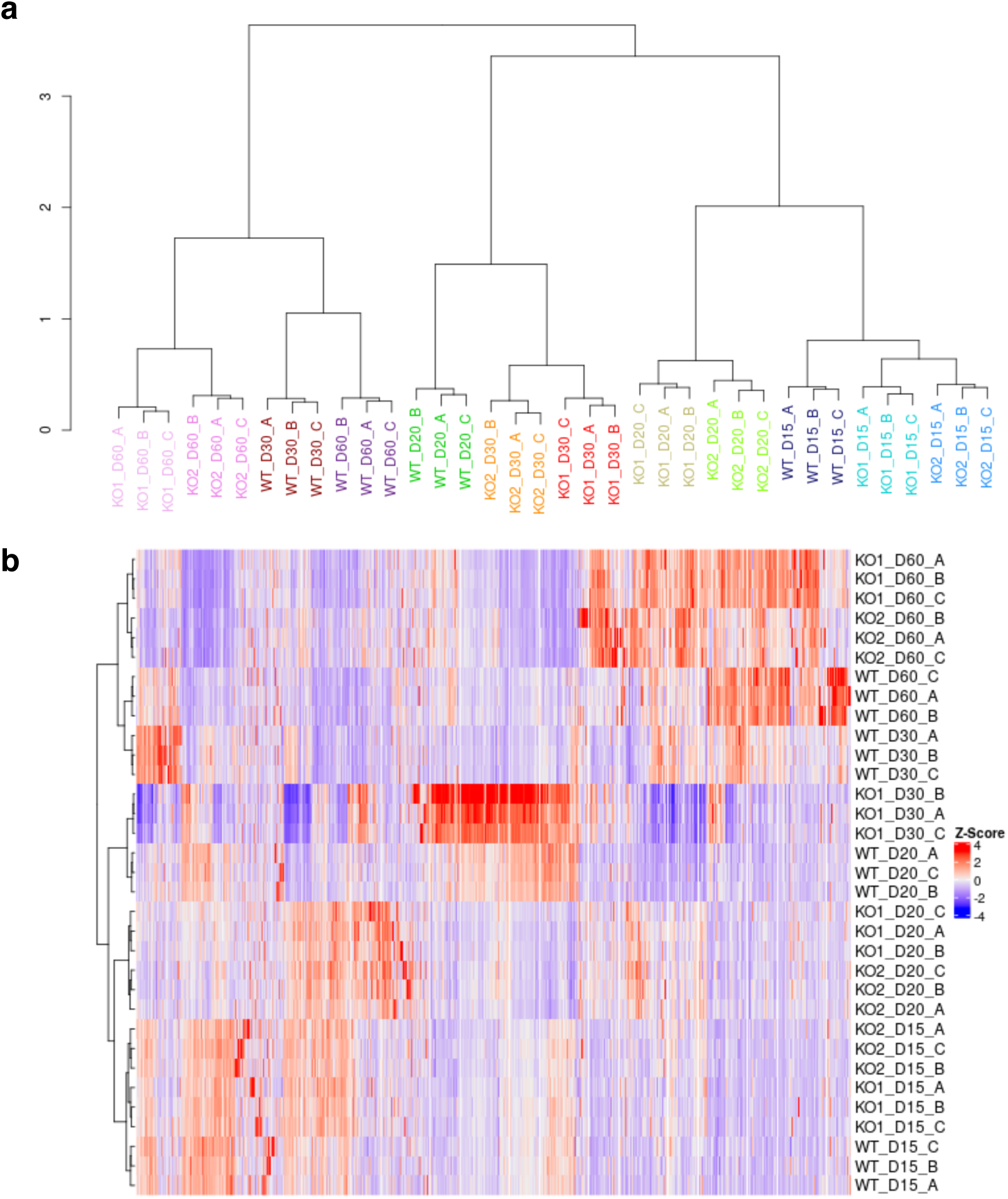
Clustering and visualisation of KO and WT sample gene expression. **a**, Hierarchical clustering of KO and WT samples based on relative expression of protein-coding genes (20,004 genes). Clustering was performed on a Pearson’s correlation matrix of the z-scored gene expression, using Ward’s linkage criterion (R hclust option ‘*ward.D2’*) and 1-r as the distance function, where r is the pairwise Pearson’s correlation coefficient. **b**, Heatmap showing transcriptomic profiles of KO and WT samples (low-quality KO2 samples from day 30 removed) based on the z-scored expression levels of variably expressed protein-coding genes (coefficient of variation > 25%; 15,003 genes). Heatmap clustering was performed as in **a**, using Pearson’s correlation matrix and Ward’s linkage criterion. Clustering and heatmap both illustrate the similarity between KO lines and their delayed development relative to WT.

**Supplementary Figure 4.**
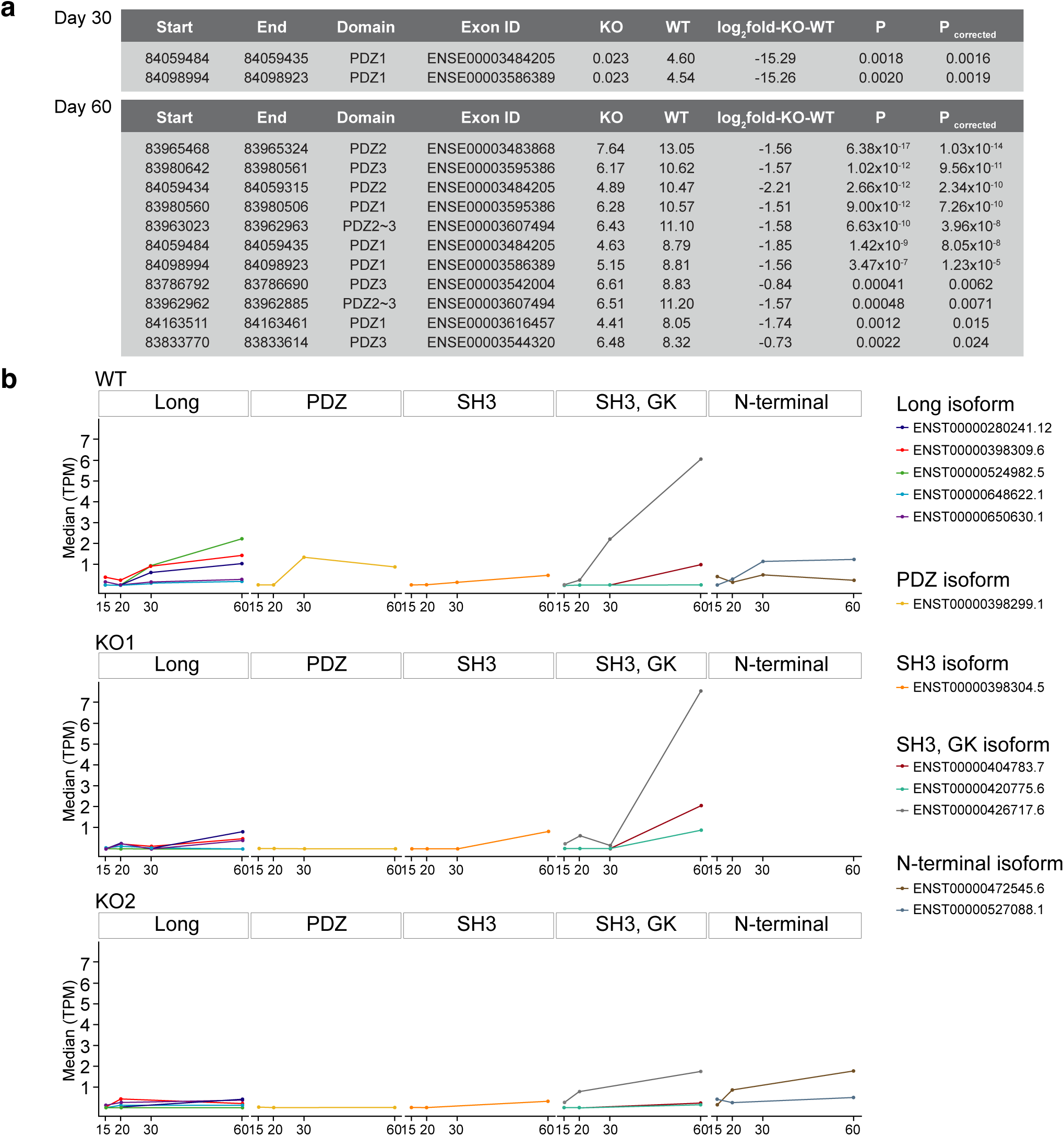
Validation of *DLG2^-/-^* human embryonic stem cells. **a**, Differential exon usage (days 30 and 60) for exons encoding *DLG2* PDZ domains. Analysis was performed using DEXseq. Shown are all exonic regions significant at an FDR of 5%. **b**, *DLG2* predicted transcript expression in WT, KO1 and KO2 lines at each timepoint. Transcript-level counts in TPM (Transcripts Per Million) were imported from RSEM using tximport (v1.12.3) (Soneson et al., 2015). Shown are all protein-coding transcripts consistently identified (TPM > 0 in all replicates) in at least one timepoint. Transcripts were classified according to the functional domains they contain and the median TPM plotted for each timpoint. Long transcripts contain all 3PDZs, SH3 and GK domains.

**Supplementary Figure 5.**
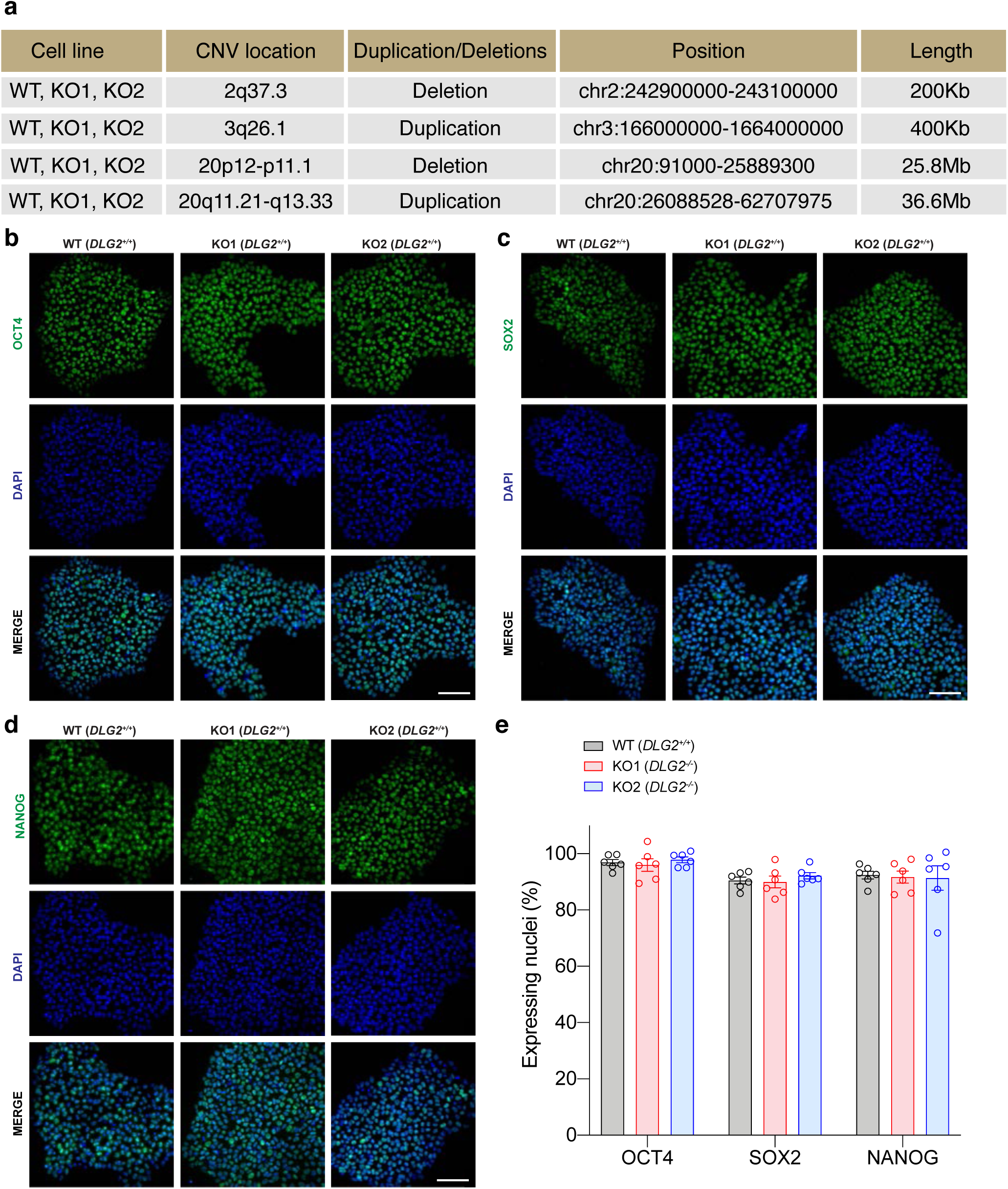
Quality control of *DLG2^-/-^* human embryonic stem cells. **a**, Copy number variant analysis of *DLG2^-/-^* hESCs. WT, KO1 and KO2 genomic DNAs were analysed using Illumina PsychArray v1.1 and data analysed using PennCNV. CNVs smaller than 100Kb and containing less than 10 SNPs were filtered out in the PennCNV QC. No additional CNVs are identified in both KO lines in comparison to WT line. **b**-**e**, Pluripotency marker expression of *DLG2^-/-^* hESCs. Key pluripotency transcription factors were expressed both in 2 *DLG2^-/-^* hESC lines and WT controls. Representative ICC images of OCT4 (**b**), SOX2 (**c**) and NANOG (**d**) expression with DAPI nuclear counterstain for 2 *DLG2^-/-^* hESC lines and WT controls. **e**, ICC quantification of nuclei expressing either OCT4, SOX2 or NANOG in 2 *DLG2^-/-^* hESC lines and WT controls, the genotype had no significant effect on the expression of these markers (OCT4: F_2,15_=0.3780, P=0.9616, n=6; SOX2: F_2,15_=0.5383, P=0.5946, n=6; NANOG: F_2,15_=0.03433, P=0.9663, n=6). Analysis was by one-way ANOVA for each pluripotency marker. All data presented as mean ± SEM and all scale bars are 100 μm.

**Supplementary Figure 6.**
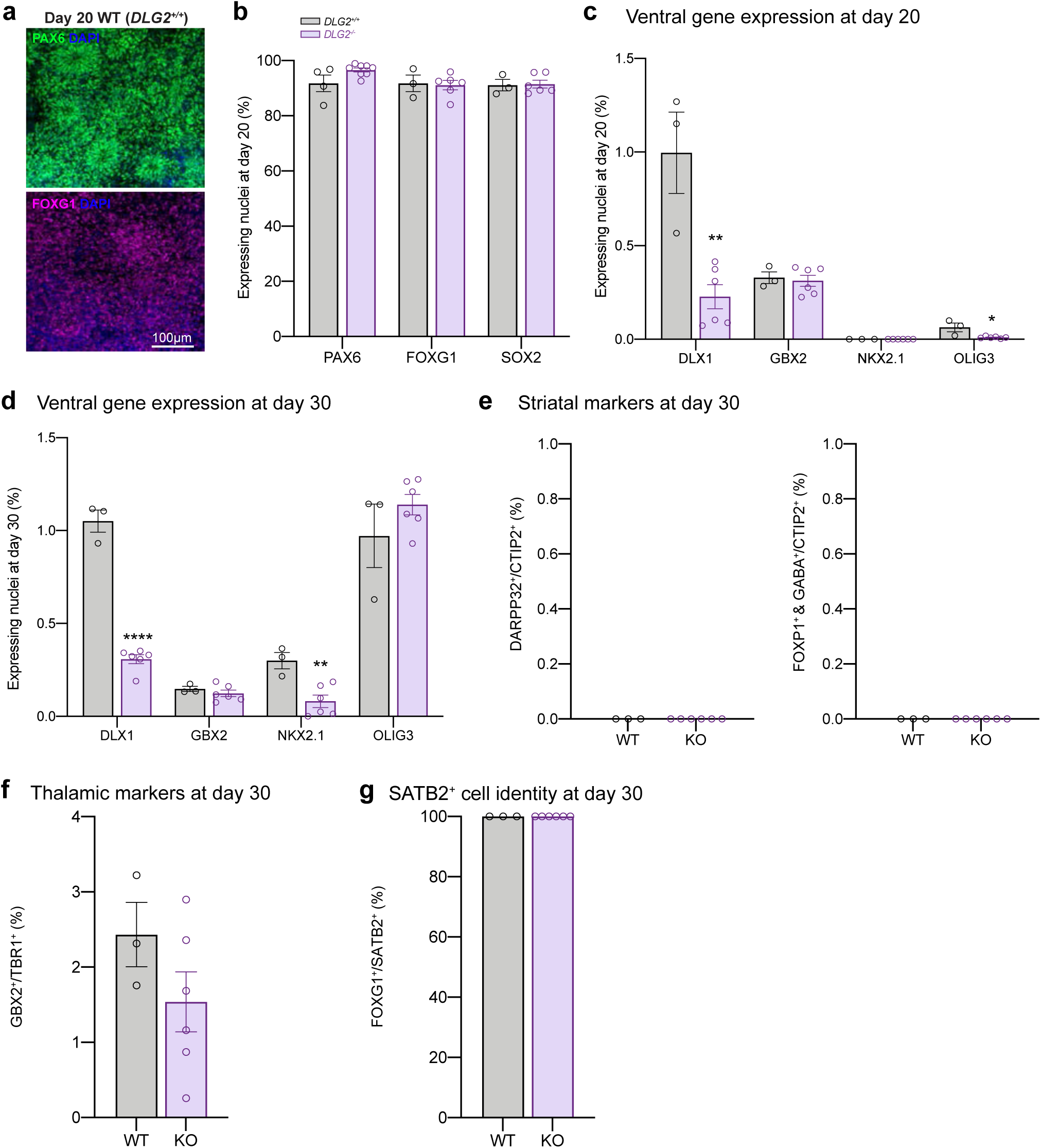
Cell fate characterisation of WT and *DLG2^-/-^* cells during cortical differentiation. **a**, Representative ICC images showing expression of forebrain progenitor markers PAX6 and FOXG1 at day 20 of cortical differentiation. **b**, ICC quantification of nuclei expressing either PAX6, FOXG1 or SOX2 in *DLG2^-/-^* and WT cells at day 20 of cortical differentiation, the genotype had no significant effect on the expression of these markers (PAX6: t_10_=2.113, P=0.0607, n≥4; FOXG1: t_7_=0.2133, P=0.8372, n≥3; SOX2: t_7_=0.1702, P=0.8697, n≥3). **c**, ICC quantification of nuclei expressing either DLX1, GBX2, NKX2.1 or OLIG3 in *DLG2^-/-^* and WT cells at day 20 of cortical differentiation. Cells epressing these markers were minor, less than 1%. DLX1: t_7_=4.521, P=0.0027, n≥3; GBX2: t_7_=0.3386, P=0.7448, n≥3; OLIG3: t_7_=3.414, P=0.0112, n≥ 3. **d**, ICC quantification of nuclei expressing either DLX1, GBX2, NKX2.1 or OLIG3 in *DLG2^-/-^* and WT cells at day 30 of cortical differentiation. Cells epressing these markers were minor. DLX1: t_7_=14.05, P<0.0001, n ≥3; GBX2: t_7_=0.8917, P=0.4021, n≥3; NKX2.1: t_7_=3.875, P=0.0061, n≥3 OLIG3: t_7_=1.219, P=0.2623, n≥3. **e**, None of the CTIP2^+^ cells co-expressed striatal markers such as DARPP32 or FOXP1 and GABA. **f**, Minority of TBR1^+^ cells expressed the thalamic marker GBX2. t_7_ =1.379, P=0.2105, n≥3. **g**, All SATB2^+^ cells expressed the telencephalic marker FOXG1. All data set except for **e** and **g** was analysed using unpaired two-tailed Student’s t-tests for each marker investigated. All data presented as mean ± SEM where possible. *, p<0.05; **, p< 0.01; ****, p<0.0001 compared to WT.

**Supplementary Figure 7.**
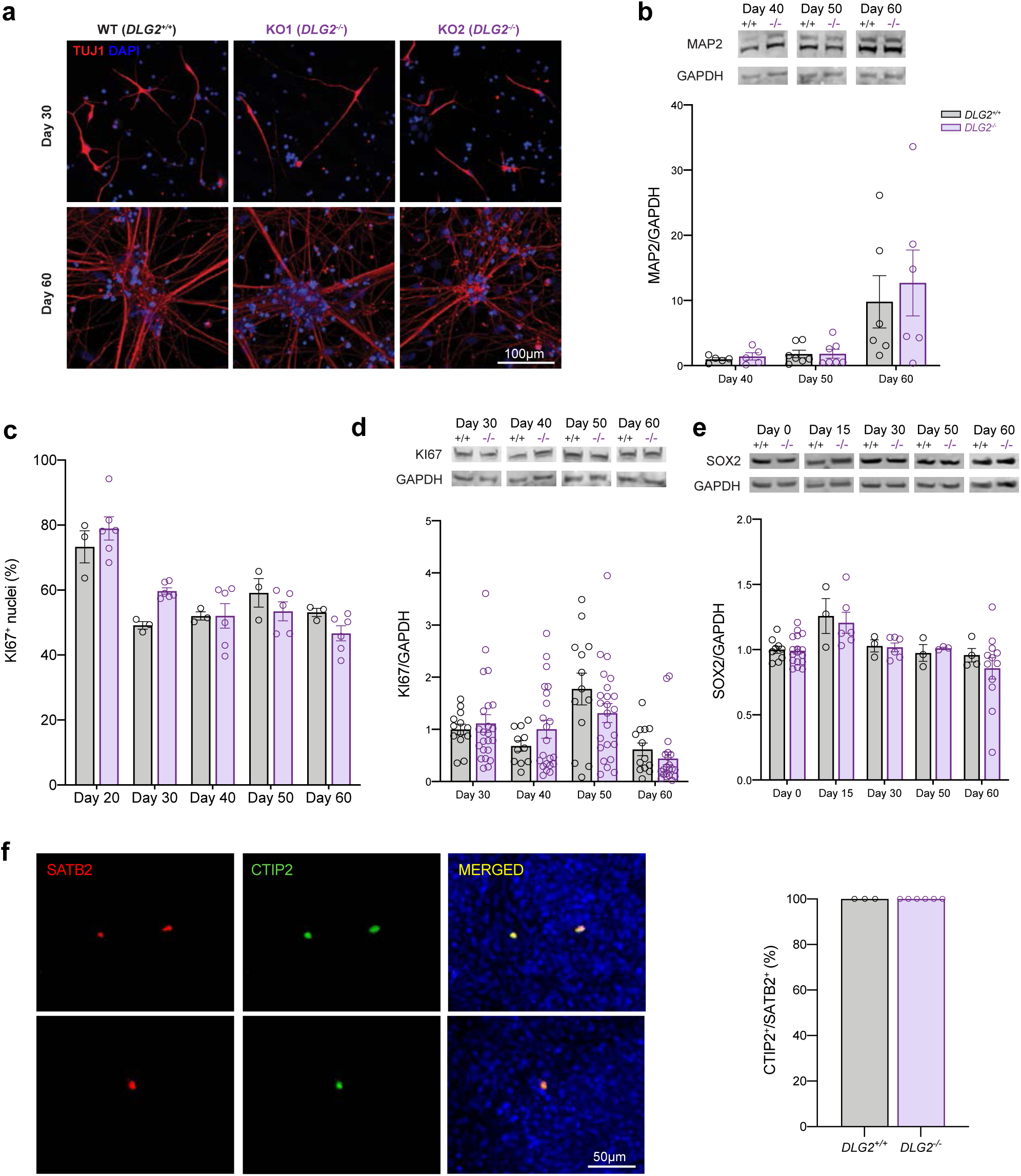
Further characterisation of WT and *DLG2^-/-^* cells during cortical differentiation. **a**, Representative ICC images showing the expression of the neuronal marker TUJ1 (1-Tubulin III) in 2 *DLG2^-/-^* lines and WT controls at days 30 and 60 of cortical differentiation. **b**, MAP2 western blot protein bands and histograms of expression normalised to GAPDH and day 40 WT expression for *DLG2^-/-^* and WT cells at 3 time points of cortical differentiation. Time (F_2,30_=8.721, P=0.0010, n≥5) did have a significant effect on MAP2 expression, while genotype (F_1,30_=0.2673, P=0.6157, n≥5) did not. **c**, ICC quantification of KI67 expressing nuclei for *DLG2^-/-^* and WT cells at 5 time points of cortical differentiation. Time (F_4,34_=20.71, P<0.0001, n≥3) did have a significant effect on KI67 expression, while genotype (F_1,34_=0.1535, P=0.6976, n≥3) did not. **d**, KI67 western blot protein bands and histograms of expression normalised to GAPDH and day 30 WT expression for *DLG2^-/-^* and WT cells at 4 time points of cortical differentiation. Time (F_3,132_=10.94, P<0.0001, n≥11) did have a significant effect on MAP2 expression, while genotype (F_1,132_=0.1536, P=0.6957, n≥11) did not. **e**, S0X2 western blot protein bands and histograms of expression normalised to GAPDH and day 0 WT expression for *DLG2^-/-^* and WT cells at 5 time points of cortical differentiation. Time (F_4,56_=4.856, P<0.0020, n≥3) did have a signifi-cant effect on S0X2 expression, while genotype (F_1,56_ =0.2928, P=0.5906, n≥3) did not. **f**, Examples of SATB2^+^ cells co-expressing CTIP adn quantification. All data sets except those in f were analysed by two-way AN0VA with post hoc comparisons using Bonferroni correction, comparing to WT controls. All data presented as mean ± SEM.

**Supplementary Figure 8.**
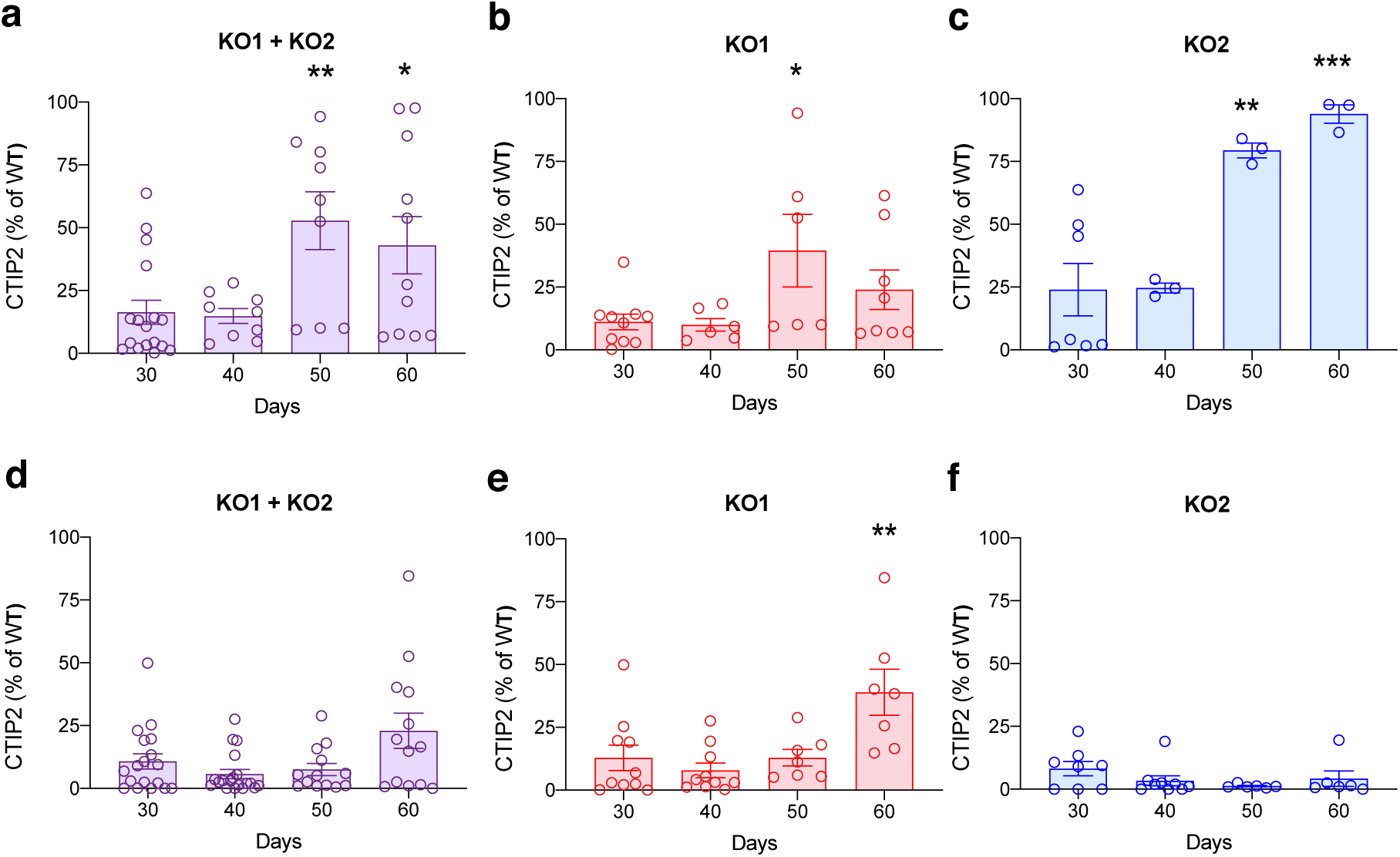
CTIP2 analysis of two *DLG2^-/-^* cell lines. **a**, The proportion of CTIP2^+^ cells in both *DLG2^-/-^* lines relative to WT level from ICC analysis. One-way ANOVA ^F^_3,42_ =5.391, P=0.0031. **b**, The proportion of CTIP2^+^ cells in KO1 *DLG2^-/-^* line relative to WT level from ICC analysis. One-way ANOVA F_3,26_=3.053, P=0.0462. **c**, The proportion of CTIP2 cells^+^ in KO2 *DLG2^-/-^* line relative to WT level from ICC analysis. One-way ANOVA F_3,12_=12.61, P=0.0005. **d**, The level of CTIP2 protein expression in both *DLG2^-/-^* lines relative to WT level from Western blot analysis. One-way ANOVA F =3.939, P=0.0125. **e**, The level of CTIP2 protein expression in KO1 *DLG2^-/-^* line relative to WT level from Western blot analysis. One-way ANOVA F_3,30_ =6.375, P=0.0018. **f**, The level of CTIP2 protein expression in KO2 *DLG2^-/-^* line relative to WT level from Western blot analysis. One-way ANOVA F_3,25_=1.493, P=0.2407. Data sets were analysed by one-way ANOVA, with post hoc comparisons using Bonferroni correction in all cases. Stars above bars represent Bonfer-roni-corrected post hoc tests, *P<0.05; **P<0.01; ***P<0.001 vs. day 30. All data presented as mean ± SEM.

**Supplementary Figure 9.**
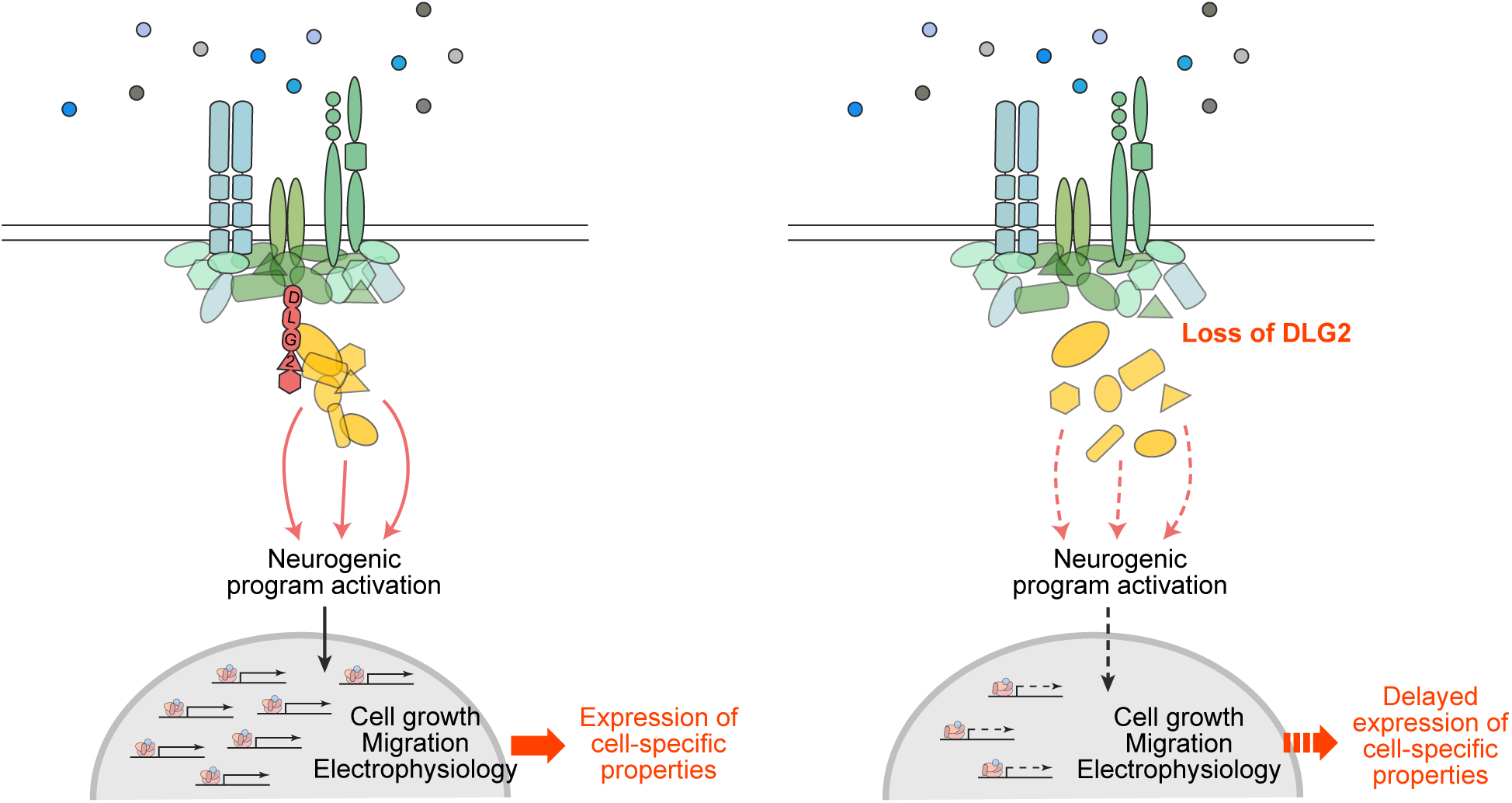
Proposed model for DLG2 action in neurodevelopment. External cues transduced by DLG2-scaffolded complexes regulate transcriptional accessibility and/or activation of neurogenic programs underlying cell growth, migration and development of electrophysiological signalling properties. DLG2 knockout impairs signal transduction, disrupting the orchestration of events required for normal development and leading to stochastic, imprecise signaling that delays expression of cell-specific properties.

Supplementary Table 1. CRISPR/Cas9 off-target validation

Supplementary Table 2. DLG2 unique peptides (LC-MS/MS of day 30 and 60 samples)

Supplementary Table 3. Differential gene expression (KO v WT and successive WT timepoints)

Supplementary Table 4. GO over-representation analysis (KO vs WT day 30 down-regulated genes)

Supplementary Table 5. GO over-representation analysis (neurogenic transcriptional programs)

Supplementary Table 6. Schizophrenia GWAS enrichment (GO terms over-represented in early-stable^-/-^)

Supplementary Table 7. GWAS enrichment in GO terms over-represented amongst early-stable^-/-^ and early-increasing^-/-^

Supplementary Table 8. *De novo* LoF enrichment in GO terms over-represented amongst early-stable^-/-^ and early-increasing^-/-^

Supplementary Table 9. Expression of neurogenic programs acrosss human *in vivo* neurodevelopmental cell-types

Supplementary Discussion

