## Supplementary Discussion for "*DLG2* knockout reveals neurogenic transcriptional programs underlying neuropsychiatric disorders and cognition"

The birth of neurons during CNS development is a sequential process involving the generation of progressively more specialised cell-types. Proliferating neuroectodermal cells (NSCs) in the neural tube give rise to neural precursors (NPCs): radial glia (RG) and intermediate progenitors (IPCs). Both RG and IPCs have a limited ability to proliferate, with RG giving rise to neurons either directly or via IPCs. In the cortex, excitatory neurons in layers VI-II are born in a sequential, inside-out fashion (from deep to upper layers), with early-born post-mitotic cells becoming deep layer (VI, V) neurons. This process is broadly recapitulated by *in vitro* differentiation protocols (e.g. Fig. 1b), with neurons produced between our day 30 and 60 timepoints predominantly displaying a deep layer identity.

*DLG2*$^{-/-}$ lines give rise to fewer neurons expressing deep layer marker CTIP2 (*BCL11B*), although bulk mRNA levels recover fully by day 60 (Supplementary Table 3). The proportion of CTIP2$^+$ cells also significantly recovers over time (Supplementary Fig. 8). Neuronal identity is thought to be determined by the internal state of NPCs immediately prior to cell-cycle exit, with changes in this state over time leading to the progressive generation of layer VI-II cells[1]. This would suggest that *DLG2* loss causes a delay in new-born neurons exhibiting their latent, deep layer identity. This also matches the observed pattern of TBR1 expression. TBR1 is expressed by virtually all post-mitotic glutamatergic neurons *in vivo* but has markedly higher expression in early-born neurons[2,3], i.e. while basal levels of TBR1 are a feature of all new-born neurons, higher levels are associated with neurons manifesting deep layer properties. The proportion of TBR1$^+$ neurons does not differ between *DLG2*$^{-/-}$ and wild-type (Fig. 2d, g), consistent with no change in the rate of neurogenesis. However, total TBR1 mRNA is significantly decreased in *DLG2*$^{-/-}$ lines at day 30 (Supplementary Table 3) indicating a reduced level of expression in individual new-born neurons (predominantly deep layer subtypes, as noted earlier). This reduction in TBR1 mRNA recovers by day 60 (Supplementary Table 3), consistent with *DLG2* loss delaying the manifestation of deep layer identity. The wider changes in gene expression and neuronal physiology seen in *DLG2*$^{-/-}$ lines are also consistent with such a delay. Loss of *DLG2* suppresses expression of many genes (85%) normally up-regulated between days 20 and 30 in wild-type, coinciding with the onset of neurogenesis. Consistent with the delayed activation of transcriptional programs underlying neurogenesis, gene expression in *DLG2*$^{-/-}$ samples from day 20 onwards resembles that of WT samples from the preceding timepoint (Supplementary Fig. 3). Prominent amongst those suppressed are genes contributing to the development of cell morphology, connectivity and electrophysiological properties (Fig. 2b, 8a) – key features of neuronal identity – leading to developing neurons with long, sparsely branched primary neurites and immature action potentials (Fig. 3, 5). Gene expression recovers somewhat by day 60 (Supplementary Fig. 3), at which point none of the above processes (morphology, connectivity, electrophysiology) are over-represented amongst genes dysregulated in *DLG2*$^{-/-}$ lines (data not shown). Morphological deficits persist at least until day 70, although the extent to which delayed upregulation of gene expression impacts mature neuronal identity is unclear: future work will need to investigate the persistence of deficits in hESC-derived *DLG2*$^{-/-}$ neurons following transplantation into rodent brain.

The commitment of precursors to a neuronal fate triggers an orchestrated sequence of events requiring the timely activation and repression of multiple transcriptional programs. Here we identify a cascade of transcriptional programs active during early corticoneurogenesis. While much further work is required to validate and refine this model it provides a useful conceptual framework that locates many genetic risk factors in a coherent sequence of cellular events, generating insight into the overlapping aetiologies of Mendelian and genetically complex neurodevelopmental disorders (Fig. 4, 6, 7, 8a).

Major changes in cell identity require alteration of chromatin structure to reflect cell-type differences in gene transcription[4]. This is evident in early-transient$^{-/-}$, in which histone- and chromatin-binding genes are highly over-represented (Supplementary Table 5, Fig. 8a). Known risk genes include chromatin modifier *CHD8* (ASD[5-7]) and histone methyltransferase *SETD1A* (NDD & SZ[8]). We hypothesise that re-organisation of the transcriptional landscape by early-transient$^{-/-}$ leads to activation of a second expression program (early-stable$^{-/-}$) which oversees a prolonged phase of cell growth and differentiation, accompanied *in vivo* by radial migration into the appropriate cortical layer (Fig. 8a). Supporting this, targets of *CHD8* are highly over-represented in early-stable$^{-/-}$ (Fig. 4d) indicating an important role in regulating the activity of this program. These targets include transcription factor *TCF4*, another major regulator of early-stable$^{-/-}$ genes (Fig. 4d) and a known interactor of pro-neural genes during differentiation[9]. Rare mutations in *TCF4* lead to Pitt-Hopkins syndrome, whose features include ID/NDD[10]; genome-wide significant (GWS) SZ association has also been identified at the *TCF4* locus[11,12]. Also present are *TRIO* (SZ[13]), which promotes reorganization of the actin cytoskeleton during neurite outgrowth[14,15], and *FMR1* (FMRP) (fragile-X syndrome[16], ASD[17], SZ[12]). While discussion of FMRP in disease has tended to focus on its role in plasticity at mature synapses[17-19], it is also present in the axons and dendrites of developing neurons where it controls growth cone morphology and motility[20]. Reflecting this role, targets of FMRP are highly over-represented in early-stable$^{-/-}$ (Fig. 4d).

*TCF4* and FMRP also regulate components of a third program, early-increasing$^{-/-}$. Functional classes of genes over-represented in early-increasing$^{-/-}$ suggest that it shapes the emergence of cell-type specific properties: dendrite morphology, synaptic connectivity and neuronal activity (Fig. 4e, 8a). This program contains transcriptional regulators *TBR1* and *BCL11B* (CTIP2), markers of deep layer neurons. *TBR1* is enriched for rare coding variants in ASD cases[21], while *BCL11B* lies in a single-gene GWS SZ locus[12,22]. Rare coding variation in genes controlling AP propagation has previously been implicated in ASD[23]; here we find that early-increasing$^{-/-}$ genes regulating action potential (AP) firing display stronger evidence for SZ association than other elements of this program (Fig. 4e). These 8 genes encode: voltage-gated sodium channels *SCN1A-SCN3A* ($Na_V$1.1-1.3) and *SCN3B* ($Na_V\beta3$); voltage-gated calcium channels *CACNA1C* ($Ca_v1.2$), *CACNA1I* ($Ca_v3.3$), *CACNA2D1 ($Ca_v\alpha2\delta$)*; and hyper-polarization activated cyclic-nucleotide-gated channel *HCN2*. $Na^+$ influx through $Na_v1.1$-1.3 drives membrane depolarisation during AP firing, with $K^+$ efflux driving repolarisation. Membrane hyperpolarisation activates HCN channels, allowing mixed $Na^+/K^+$ currents which help restore the membrane potential to its resting level[24]. T-type ($Ca_v3.3$) and L-type ($Ca_v1.2$) calcium channels contribute to neuronal firing by initiating rebound bursting and controlling calcium-sensitive potassium channels respectively[25]. Mutations in $Na_v1.2$ (*SCN2A*) lead to a wide spectrum of epileptic disorders, ID and ASD[26]. An excess of rare variants in $Na_v$ genes has also recently been reported for SZ[27]. Mutations in $Ca_v1.2$ (*CACNA1C*) lead to Timothy syndrome[28] and Brugada syndrome[29]; both *CACNA1C* and *CACNA1I* are located in single-gene GWS SZ loci[12]. Intriguingly, enhancement of spontaneous calcium transients by $Na_v$ activity impedes migration and induces premature dendritic branching in developing neurons[30]. $Ca_v1.2$ (*CACNA1C*) plays a critical role in spontaneous calcium transient generation in the axons and dendrites of developing cortical neurons, with loss of *CACNA1C* reducing neurite growth and a Timothy syndrome gain-of-function mutation impairing radial migration[31]. This suggests that, in addition to their role in AP firing, early-increasing$^{-/-}$ $Na_v$ and $Ca_v$ expression may drive an increase in calcium transients that acts as a cue for neurons to end migration and initiate terminal differentiation (dendritic branching and synaptogenesis). Deficits in this process could thus alter cell-type specific morphology and connectivity, perturbing network formation. Consequently, reduced $Na_v$ and $Ca_v$ gene expression (Supplementary Table 3) may contribute to the reduced neurite branching seen in developing *DLG2*$^{-/-}$ neurons.
